# Neurophysiological predictors of memory impairment in schizophrenia: decoupled slow-wave dynamics during non-REM sleep

**DOI:** 10.1101/578039

**Authors:** Ullrich Bartsch, Andrew Simpkin, Charmaine Demanuele, Erin Wamsley, Hugh Marston, Matthew W Jones

## Abstract

The slow-waves (SW) of non-rapid eye movement sleep (NREM) reflect neocortical components of network activity during sleep-dependent information processing; their disruption may therefore contribute to impaired memory consolidation. Here, we quantify SW dynamics relative to motor sequence memory in patients suffering schizophrenia and healthy volunteers.

Patients showed normal intrinsic SW properties but impaired SW coherence, which failed to exhibit the learning-dependent increases evident in healthy volunteers. SW-spindle phase amplitude coupling across distributed EEG electrodes was also dissociated from experience in patients, with long-range fronto-parietal and -occipital networks most severely affected. Partial least squares regression modelling confirmed distributed SW coherence and SW-spindle coordination as predictors of overnight memory consolidation in healthy controls, but not in patients.

Quantifying the full repertoire of NREM EEG oscillations and their long-range covariance therefore presents learning-dependent changes in distributed SW and spindle coordination as fingerprints of impaired cognition in schizophrenia.

## Introduction

NREM sleep EEG features two cardinal oscillations, which co-occur in systematically varying proportions during the night: deep NREM sleep (stage N3, or ‘slow wave sleep’) is dominated by low-frequency power (0.5-4Hz, sometimes subdivided into SW at 0.5-1.5Hz and delta oscillations at 1.5-4Hz) whereas lighter NREM (stage N2) encompasses the majority of 12-15Hz spindle oscillations. SWs and spindles are cortical signatures of the patterned thalamocortical and limbic network activity that supports NREM sleep’s roles in overnight memory consolidation (Plihal and Born, 1997; Walker and Stickgold, 2004; Antony et al., 2012; Cairney et al. 2018). For example, regional increases in SW density and amplitude correlate with declarative memory consolidation (Huber et al., 2004) and augmenting SW activity during sleep can enhance memory (Marshall et al., 2006; Ngo et al., 2013; Binder et al., 2014). The duration of stage N2 sleep and spindle density also correlate with performance in both procedural and declarative memory tasks (Gais et al., 2002; Walker et al., 2002; Schabus et al., 2004; Clemens et al., 2005; Barakat et al., 2011).

It follows that memory impairments in disease may be caused or exacerbated by disrupted NREM neurophysiology. This link is best characterized in schizophrenia (Manoach and Stickgold, 2009), where deficits in sleep-dependent memory consolidation correlate with aberrant EEG signatures during NREM sleep (Ferrarelli et al., 2010; Keshavan et al., 2011; Ramakrishnan et al., 2012). Some studies report that deep NREM sleep is reduced in patients (Monti and Monti, 2004; Sekimoto et al., 2007; Sarkar et al., 2010), and attenuated SW power (Keshavan et al., 1998; Hoffmann et al., 2000; Göder et al., 2003) has also been linked to cognitive deficits (Ganguli et al., 1987; Göder et al., 2006; Ramakrishnan et al., 2012). However, not all studies report SW abnormalities (Ferrarelli et al., 2010). Schizophrenia has more consistently been associated with reductions in spindle density or sigma power in both patients and first-degree relatives (Ferrarelli et al., 2007, 2010; Manoach et al., 2010; Seeck-Hirschner et al., 2010; Keshavan et al., 2011; D’Agostino et al., 2018), with spindle measures showing some correlation with impaired sleep dependent memory consolidation in patients (Wamsley et al., 2012).

Paralleling the emergence of this clinical evidence linking disrupted sleep-dependent neural network activity with impaired memory consolidation, deep-brain recordings in rodents have detailed the roles of NREM network oscillations in shaping the patterned neural activity underpinning mnemonic processing. Much of this rodent work has focused on the hippocampus, where pyramidal neuron spiking patterns during NREM “replay” sequential activity encoding recent behavioral experience (O’Neill et al., 2010). However, the timing of hippocampal activity during NREM is influenced by SW activity (Isomura et al., 2006; Hahn et al., 2012; Taxidis et al., 2013) and is in turn coordinated with spindles in neocortex (Siapas and Wilson, 1998; Sirota et al., 2003; Mölle et al., 2006; Clemens et al., 2007; Wierzynski et al., 2009). Beyond examining SW or spindles *per se*, quantifying the coordination between ripples, spindles and SW can therefore illuminate the mechanisms and functions of brain activity during sleep in health and disease (Gardner et al., 2014; Latchoumane et al., 2017).

(Phillips et al., 2012) began to address disordered interdependence of SW, spindles and ripples using the Methylazoxymethanol acetate (MAM)-E17 rat neurodevelopmental model of schizophrenia (Moore et al., 2006), demonstrating that the MAM-E17 model harbors reduced NREM SW power and reduced coherence of SW between frontal and occipital cortices. This reduced SW coherence was accompanied by impaired frontal slow-wave phase coupling to posterior cortical spindle amplitude and decoupling of cortical spindle and hippocampal ripple oscillations, potentially as a consequence of interneuronal dysfunction (Phillips et al., 2012). Consistent with predictions from this rodent work, a recent analysis by Demanuele et al. showed that regression models incorporating both spindle density and a measure of spindle-SW phase-locking were able to predict memory performance in schizophrenia patients (Demanuele et al., 2017). However, Demanuele et al. focused on local SW-spindle phase-locking (SW and spindles detected on the same EEG recording site), which did not predict sleep-dependent memory in healthy participants.

These results prompted us to investigate properties of SWs and their interrelationship with spindles across different cortical sites using the same sleep EEG recordings from chronic medicated schizophrenia patients and demographically-matched control participants as Demanuele et al. (2017) and (Wamsley et al., 2012). Wamsley et al. reported reduced spindle density and 12-13.5Hz (low sigma) power during N2 sleep in these patients, alongside reduced spindle coherence between centro-occiptal EEG electrodes. Of these measures, spindle densities correlated with overnight improvement on the finger tapping motor sequence task (MST, Karni et al., 1998; Walker et al., 2002), but only in schizophrenia patients and not in healthy volunteers. Here, we show that quantification of SW properties, SW coherence, and SW-spindle phase-amplitude coupling across the cortical mantle strengthens and clarifies links between NREM neurophysiology and memory consolidation. Our results highlight the importance of long-range network connectivity during NREM as central to mnemonic processing in healthy volunteers, and dissociation of SW coherence from experience in patients diagnosed with schizophrenia.

## Results

### Impaired sleep-dependent motor memory consolidation in SCZ

We present novel analyses of a previously acquired dataset (Wamsley et al., 2012). Briefly, participants were invited to the sleep lab for a baseline polysomnography (PSG) recording on night 1 (‘base’). Night 2, which we refer to as the learning night (‘learn’, Fig. 1a), was flanked by a motor sequencing task before (MST train) and after (MST test) sleep. For the current analysis of behavior we only included participants with quality-controlled EEG recordings on night 2 (see Methods). Patients diagnosed with schizophrenia (SCZ) and healthy controls (CON) showed similar normalized learning curves during training on the MST. However, patients did not show overnight improvement on the MST compared to healthy controls (improvement percentages CON mean 15.93, sd = 13.68; SCZ: mean 1.62, sd=22.88, two-sided t-test, p = 0.025, t-stat: 2.34 df: 33.17, Figure 1b, see Wamsley et al., 2012 for results from a larger sample).

**Figure 1:**
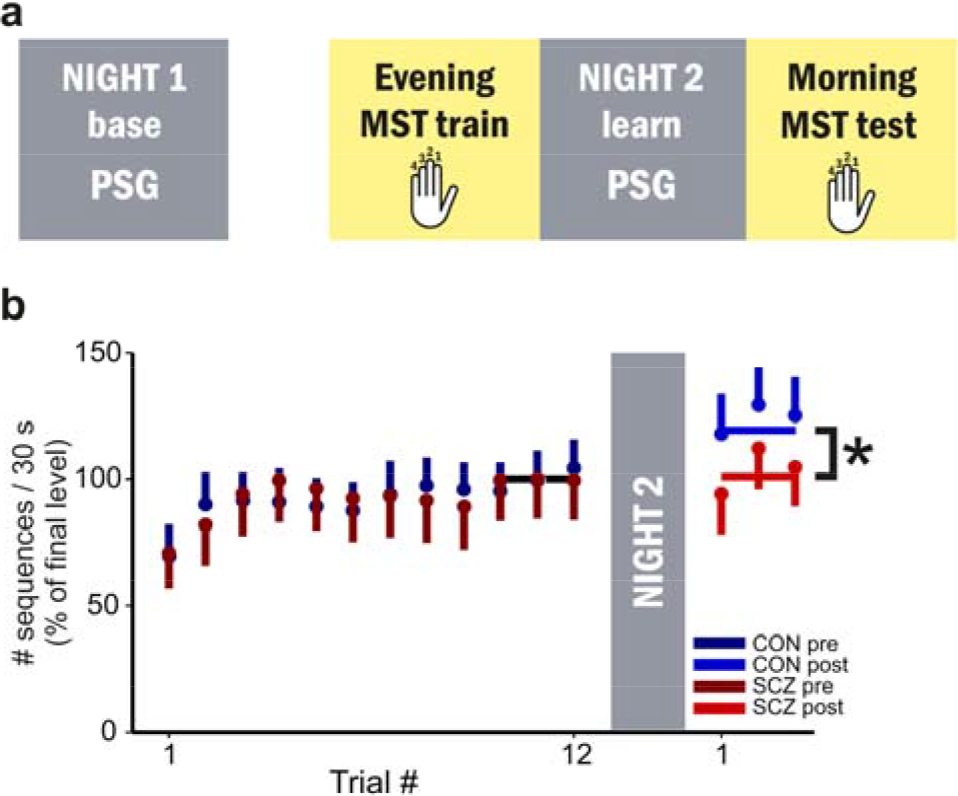
Patients with schizophrenia show impaired sleep-dependent memory consolidation in the motor sequence task (MST). **a) Timeline of behavioral testing and polysomnography.** The baseline night 1 (base) was not preceded by any behavioral training. The following night, participants were trained on the MST 1h before their regular bed time (MST train) and were tested the following morning, after up to 10h sleep during night 2 (learn). **b) MST performance.** Behavioral results for all participants included in night 2 EEG analyses (healthy controls, CON, n=15; patients, SCZ, n=21). Because SCZ patients typed fewer correct sequences during training, task performance (number of correct sequences/30s) was normalized by the asymptotic level achieved during MST training (mean of last 3 trials, <CON_10-12_>=19.05, <SCZ_10.12_>= 11.52). Controls show a significant improvement on the MST on the next day, whereas SCZ do not (improvement in percentage, two-sided t-test, p = 0.025, t= 2.3,4 df: 33.17, see Wamsley et al., 2012, for statistics on a larger sample).

### Conserved slow-wave event properties in SCZ

*We* analyzed artefact-free EEG traces from all epochs of N2 and N3 sleep (referred to collectively as NREM sleep) at the electrode positions depicted in Figure 2a. Comparisons of the basic spectral properties of all NREM sleep epochs show that the topographic distributions of SW and spindle power were similar in volunteers and patients (Supplementary Figure S1a, b), though spectra at individual electrodes showed increased power at sub-10Hz frequencies in patients, particularly over central and occipital electrodes (Supplementary Figure S1c).

**Figure 2:**
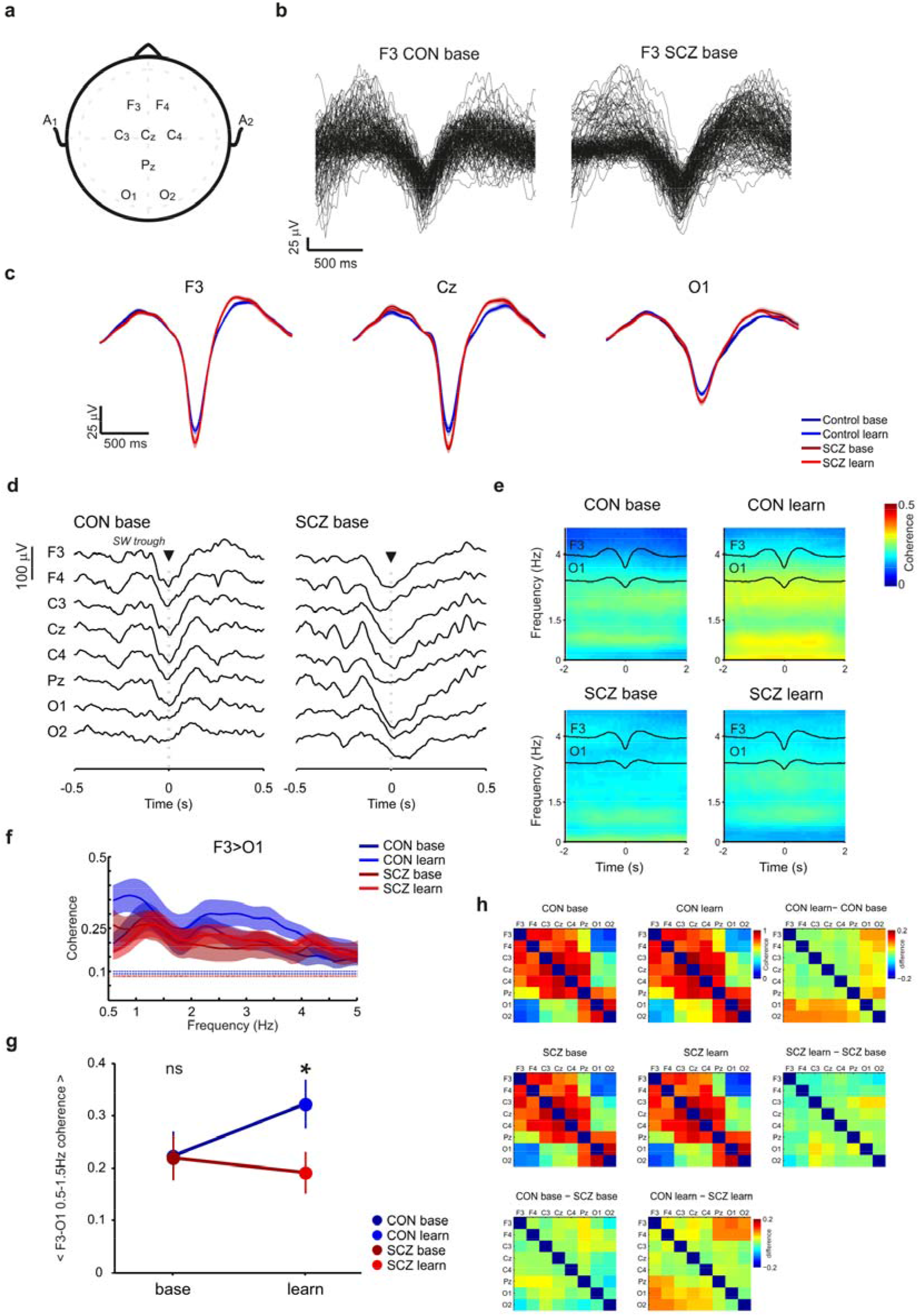
Normal SW amplitudes but reduced learning-dependent SW coherence in SCZ during NREM sleep. **a) EEG recording configuration.** EEG was recorded from F3, F4, C3, Cz, C4, Pz, O1, O2, but some electrodes were lost during recording, so n numbers vary for different electrode positions (all recordings: CON base and learn n=15; SCZ base, n=19, SCZ learn= 21). **b) Example slow waves (SW).** Overlays of all automatically detected slow waves in wideband EEG from one healthy control (CON) and one patient (SCZ) during baseline NREM sleep at electrode F3. **c) Average wave triggered averages of detected SW at F3, Cz and 01.** Wave triggered SW averages in controls (dark blue: baseline, light blue: learning) and patients (dark red: baseline, light red: learning). Patients show a slight increase in absolute amplitude of detected SW events most prominent nearthe peaks of the SW (UP states), although significant bins did not survive Benjamini-Hochberg correction (FDR). See Supplementary Figure S2b for uncorrected significant differences between CON and SCZ) **d) Example EEG traces during SW at F3 in CON and SCZ.** Examples of an automatically detected SW at F3 and 1 second windows of simultaneously recorded activity across all other electrodes during the baseline night. EEG traces are centered on the SW trough time at F3. **e) Average SW triggered multi-tapered coherograms between F3 and O1 during NREM sleep.** All trough times from SWs detected at F3 were used to select a ±2s window at F3 and 01 to compute a sliding window multi-taper coherogram. Wave triggered averages of SWs at the trigger channel (F3, top trace) and target channel (01, bottom trace) are overlaid. Coherograms (included F3-O1 pairs: N_CON_=11, N_SCZ_=14) were averaged for each recording night and group. CON subjects show an increase in coherence around SWs following MST learning, whereas SCZ patients do not. **f) Average SW triggered coherency during NREM sleep** All peak times from SWs detected at F3 were used to compute the average coherency between F3 and O1 in a ±2s window. Participant coherency (included F3-O1 pairs: N_CON_=H, N_SCZ_=14) was averaged for each recording night and group and jackknife confidence intervals were calculated. Note the learning dependent increase in the CON group in the range of 0.5-1.5 Hz. **g) Interaction plot for average SW triggered slow (0.5-1.5 Hz) coherence** Average 0.5-1.5Hz coherence values for each participant were analyzed using a linear mixed model. Least squares mean estimates of the group means were plotted in an interaction plot. A significant interaction effect between night and group emerged in the final model: Average slow coherence increases significantly after learning only in the CON group and was thus higher than in SCZ during the learning night (* p<0.05; see main text and supplementary statistical methods M2 for mixed model statistics). **h) Group differences in SW triggered slow coherence** Coherence matrices display means of SW coherence and differences between all sensor pairs (N numbers vary for different pairings due to electrode loss in some cases: CON 8-15; SCZ 11-21). Mean coherence values (0.5-1.5 Hz, ±2s around SW) were plotted against electrode position to create a coherence matrix. Average coherence matrices were calculated for baseline (left), for learning (middle), in each group (CON, top; SCZ, middle), respectively. Differences in SW coherence between baseline and learning night for both groups are shown to the right and below the respective matrices of mean values. Note the learning dependent increases in the CON group (CON learn – CON base) in comparison to little change in the patient group (SCZ learn – SCZ base).

Next we detected individual SW events on all EEG channels as described in Phillips et al., 2012 (see also Supplementary Figure S2a). Figure 2b shows typical SWs detected at electrode F3 in one healthy participant and one SCZ patient during the baseline night. There were no differences in SW density, oscillation frequency, amplitude or duration between controls and patients (Supplementary Figures S2 and S3). To quantify SW morphology, we calculated wave-triggered averages (triggered by SW trough times) for each electrode, group, and night. Figure 2c shows average SW for F3, Cz and O1 electrodes; although SW tended to be slightly higher amplitude in patients, these differences were not significant (two-sided Wilcoxon ranksum test with FDR correction, P_BH_<0.05; see Supplementary Figure S2b for bin-by-bin analysis).

### Reduced long range coupling of 0.5-1.5Hz slow waves in SCZ

SW tend to originate in frontal cortices and can be either local or travelling waves that, in some instances, are coordinated across multiple cortical areas (as exemplified in Figure 2d; Amzica and Steriade, 1995; Achermann and Borbély, 1998; Mölle et al., 2004; Stroh et al., 2013; Miyamoto et al., 2016). To quantify coordination of distributed SW activity across EEG electrodes, we used SW troughs as reference timestamps around which to calculate multi-tapered spectral coherence (Bokil et al., 2010).

Figure 2e shows group-averaged coherograms for volunteers and patients during baseline and post-learning NREM sleep. 0.5-1.5 Hz SW coherence between electrodes F3 and O1 did not differ between CON and SCZ during the baseline night but showed a marked learning-dependent increase in CON (also shown averaged over NREM in Figure 2f) that was missing in SCZ patients. A linear mixed model analysis of Fisher z-transformed average coherence values revealed a fixed effect for recording night, although this main effect did not survive type III Analysis of Variance with Satterwaithe approximation for degrees of freedom (F_(1,20.63)_ = 1.56, p=0.23). However, a significant night x group interaction for SW coherence was evident (F_(1,20.63)_ = 5.04, p=0.036). Thus, SW coherence increased significantly following MST training in the CON group (ls-means estimation for post-hoc testing, p_(CON base – CON learn)_ = 0.03), and was significantly higher compared to SCZ during the learning night (p_(SCZ learn – CON team)_ = 0.04, see also Supplementary Statistics Methods M1).

To visualize the topography of these group- and learning-dependent changes, average SW coherence values across all electrode pairs are shown for each group and night in Figure 2h. Following MST training, the CON group exhibited increases in fronto/central to occipital coherence (Figure 2h, CON learn – CON base) that were not evident in patients. Consequently, comparison between CON and patient SW coherence during the learning night (Figure 2h, CON learn – SCZ learn) highlights deficits in learning-dependent, long-range coordination of SW activity between frontal/central and occipital cortices in SCZ.

Overall, these SW analyses show that SCZ patients exhibit conserved SW densities and amplitudes but an absence of learning-dependent coordination of SW activity across the cortical mantle.

### Reduced learning dependent coupling of SW and spindles in SCZ

The coordination of SW with spindle oscillations has been shown to correlate with memory consolidation (Clemens et al., 2007; Cox et al., 2012, 2014) and can be readily identified in EEG recordings (see examples in Figure 3a). De Manuele et al. (2017) demonstrated that *local* slow wave phase to spindle amplitude coupling (i.e. derived from signals from the same EEG sensor) can correlate with sleep-dependent memory consolidation, but only in SCZ patients. To test whether *distributed*, cortex-wide coupling of SW and spindle oscillations is a more sensitive metric of memory consolidation and impairment, we used an established phase-amplitude coupling measure (modulation index, MI; Tort et al., 2010; Onslow et al., 2011) between the most distal electrode locations, F3 and O1. F3 SW were entered as modulating signal (phase) and target electrode O1 was entered as the modulated signal (amplitude).

**Figure 3).**
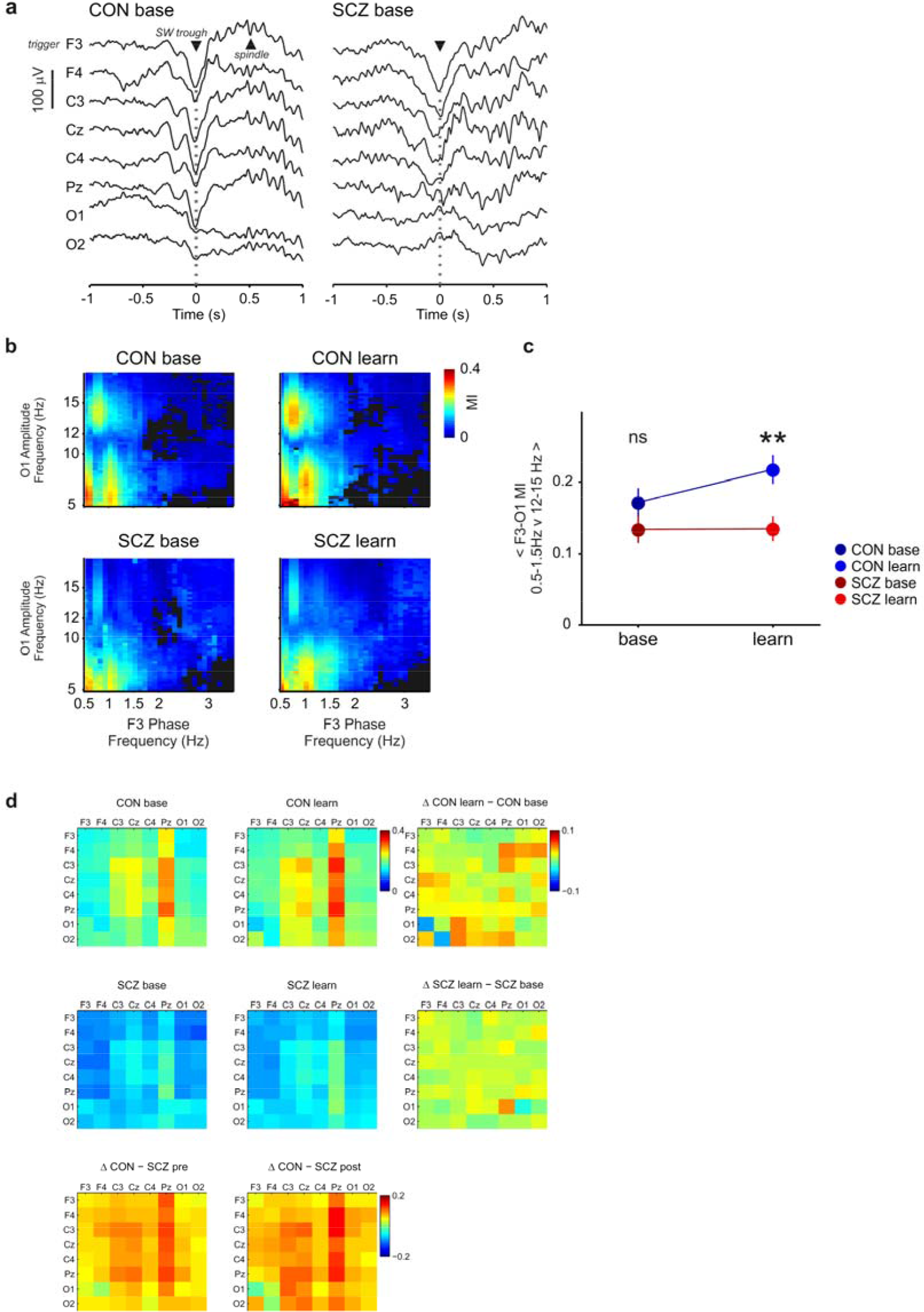
Long range SW-spindle coupling shows a learning dependent increase in CON but not in SCZ. **a) SW spindle co-occurrence during NREM sleep.** Examples of a frontally detected SW and subsequent spindle oscillations across multiple channels from one healthy CON individual (CON) and one patient (SCZ) during baseline sleep. **b) SW triggered frontal occipital phase amplitude coupling** The negative peaks of F3-derived SW were used as reference points to calculate a comodulogram for a ±2s window of data at F3 and O1. The F3 trace was used as modulating input (phase) and O1 was used as modulated input (amplitude). Only significant MI values were included in further analysis (n.s. are dark grey bins, see Methods). The resulting participant co-modulograms were averaged across nights for each group. In the CON group (left) there is a cluster of high MI values for SW phase (0.5-1.5 Hz) and spindle amplitude (12-15 Hz), this cluster of high MI values appears to increase in CON after learning. The same cluster is present in the SCZ but MI values appear smaller in comparison to CON and there is little learning dependent change. **c) Interaction plot of mean F3-O1 SW-spindle MI** The mean SW-spindle MI values (0.5-1.5Hz phase, and 12-15 Hz amplitude) were fed in to mixed linear model. The interaction plot demonstrates the significant interaction effect between night and group in the final model. Average MI values increase significantly after learning only in the CON group (see main text for details). **d) Brain wide differences in SW (0.5-1.5 Hz) spindle (12-15 Hz) MI** MI matrices display mean values of SW-spindle PAC and differences between all sensor pairs. Mean MI values (0.5-1.5Hz phase, and 12-15 Hz amplitude) were plotted against electrode position to create a PAC matrix. Average PAC matrices were calculated for baseline (left) and learning (middle) nights, in each group (CON, top; SCZ, middle), respectively. Differences in SW-spindle PAC between baseline and learning night for both groups were added to the right and below the respective matrices of mean values. Note the learning dependent changes in the CON group (CON learn – CON base).

Figure 3b shows the resulting average co-modulograms for CON and SCZ groups on both recording nights. Linear mixed model analysis of the average 0.5-1.5 Hz *vs*. 12-15Hz MI revealed main effects for group (Type III analysis with Satterwaithe approximation for degrees of freedom F_(1,22.540)_=5.78, p=0.025) and recording night (F_(1,18.626)_=5.30, p=0.033) with a significant group x night interaction (F_(1,18.626)_=4.72, p=0.043). Post hoc testing confirmed a learning dependent increase in F3-01 SW-spindle MI in CON group (LSmeans estimation, P_(CON baseline - CON learning)_ = 0.007) that was significantly higher compared to SCZ during the learning night p_(SCZ learning – CON learning)_ = 0.004, Figure 3c). Consistent with Demanuele et al. (2017), this effect was not apparent for local (within electrode) SW-spindle PAC, where MI showed a group effect but no significant interaction between group and night (Supplementary Statistical Methods M4).

Since the interrelationships between SW and spindle timing across different cortical regions proved more sensitive to learning than intra-regional SW-spindle coupling, we calculated SW-spindle PAC for all electrode pairs. 0.5-1.5 Hz SW phase at one electrode (modulating phase) was used to calculate the Modulation Index of 12-15Hz spindle power (modulated amplitude) at each other electrode; the resulting MI matrices are shown in Figure 3d. In CON, the most prominent SW-spindle coupling during pre-learning NREM sleep was between Pz and frontal-central electrodes; this coupling increased after learning, particularly for Pz-F4 and more posterior, occipital electrode pairs. Overall SW-spindle coupling appeared markedly lower in SCZ patients, most notably for Pz-frontal and -central electrodes (see CON-SZ difference matrices in Figure 3d). In contrast to CON, patient SW-spindle coupling appeared insensitive to learning, remaining very similar across baseline and post-learning nights.

### Measures of SW dynamics predict memory consolidation more accurately in controls than patients

To establish whether variables that describe SW, spindles and their coupling predict sleep-dependent changes in MST performance, we built regression models for each variable set and group, enabling unbiased detection of which EEG features best predict behavioural change. A given variable set contained values of that measure for all electrodes/pairs; we then regressed NREM sleep measures during the learning night for each variable set (for example, SW coherence) against the percentage overnight improvement in MST (based on number of correct sequences).

Partial least squares regression (PLSR) is well-suited to this task; it is similar to principal component regression and is very tolerant of high co-linearity in predictor variables (Geladi and Kowalski, 1986; Krishnan et al., 2011). The prediction error of overnight MST improvement from each model reflects the explanatory power of that variable, serving here to quantify the contributions of SW coherence and SW spindle PAC to sleep dependent memory consolidation of MST performance. The larger the prediction error, the less accurately that variable predicts behavioural change; smaller Residual Sum of Squares (RESS) values equate to better prediction, with RESS=0 meaning perfect prediction. Thus, a difference in prediction error between patients and controls may indicate a functional NREM deficit in patients. Models for NREM event properties, SW and spindles, and other variables quantifying the long-range interactions of SW and spindles (e.g. SW associated spindle power and SW associated spindle coherence) are presented in Supplementary Information (Supplementary Table S1).

Figure 4 shows final prediction error results for SW coherence and SW-spindle PAC PLSR components. SW coherence is a better predictor of MST improvement in controls compared to patients (CON: RESS= 0.96, R2 =0.86, SCZ: RESS = 4.75, R2= 0.63). Long range SW-spindle PAC is overall the best predictor in controls (Figure 4 c, CON: RESS= 0.34, R2 = 0.94) compared to all other NREM sleep variables (Supplementary Table S1). SW spindle PAC in patients is a worse predictor of percentage improvement than in controls (SCZ: RESS = 2.99, R2= 0.75). Permutation tests with 10,000 permutations of group memberships indicate that group differences in RESS are significant (SW coherence, RESS_CON-SCZ_, p = 2.0e-4, SW PAC, RESS_CON-SCZ_, p = 5.0e-4) at the Bonferroni corrected alpha level of 3.3e-3 (Figure 4 b, d, Supplementary Table S1).

**Figure 4).**
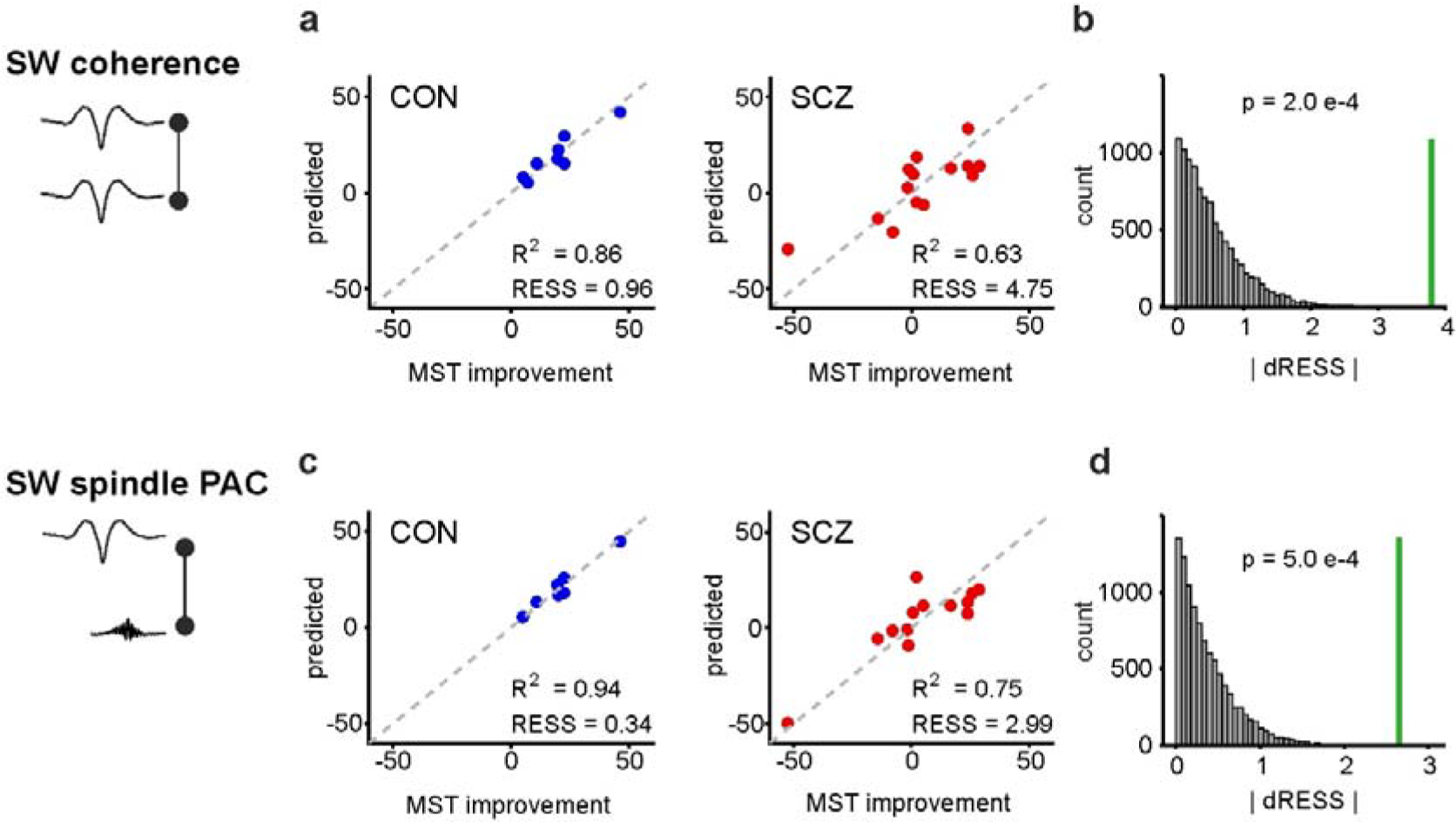
SW coherence and SW-spindle PAC predict sleep dependent memory consolidation more accurately in CON than in SCZ. **a) PLSR prediction of MST improvement from SW-triggered slow coherence** The PLSR models show that SW coherence is a better predictor (i.e. shows smaller prediction error) of overnight MST change in CON (blue symbols, Residual Sum of Squares, RESS=0.96, R^2^=0.86 using 3 components) compared to SCZ (red symbols, RESS= 4.75, R2= 0.63). **b) Permutation test result for |dRESS_CON-SCZ_| using SW triggered slow coherence as predictor** The absolute sample difference in RESS (|dRESS| = 3.79, green line) is significantly different between CON and SCZ groups (p = 2.0e-4 permutation test, n=10000, see also Supplementary Table S1, alpha = 0.0033). **c) PLSR prediction of MST improvement from SW-spindle PAC** The PLSR model for SW spindle PAC shows the smallest prediction error for MST change in CON (blue symbols, RESS= 0.34, R2= 0.94), but significantly larger prediction error for MST change in SCZ (red symbols, RESS= 2.99, R2= 0.75). **d) Permutation test result for |dRESS_CON-SCZ_| using SW-spindle PAC as predictor** The absolute sample difference in RESS (|dRESS | = 2.65, green line) is significantly different between CON and SCZ groups (p = 5.0e-4 permutation test, n=10000, see also Supplementary Table SI, alpha = 0.0033).

This PLSR analysis confirms and extends the results of linear mixed model analyses of individual electrode pair slow coherence and SW-spindle PAC, that show a specific learning dependent increases in both measures only in the control group (Figures 2–3). In conclusion, the extents of SW-SW and SW-spindle coupling are accurate predictors of sleep-dependent memory consolidation in healthy controls, but less predictive in patients.

## Discussion

We analyzed NREM sleep EEG data from patients diagnosed with SCZ and healthy controls, translating predictions arising from a rat model of impaired neurodevelopment that showed dys-coordination of NREM slow-waves and spindles across the cortical mantle (Phillips et al., 2012a). We found striking similarities between patient and rodent EEG: impaired long-range SW coordination, in particular between frontal and occipital cortices, and a disrupted nesting of centro-parietal spindles in frontal SW. These impairments in patients were most prominent following training in a motor sequence task, since NREM EEG in SCZ patients failed to show the experience-dependent increases in coordination evident in controls. Quantifying the network coordination of SW and spindles during NREM therefore generates objective measures that predict sleep-dependent memory consolidation in healthy participants, and impaired sleep-dependent memory in SCZ patients.

### Reduced long-range SW coherence in SCZ

There is mixed evidence for SW/delta deficits in patients with SCZ: a meta-analysis (Chouinard et al., 2004) failed to confirm reports of altered spectral power in the SW/delta range (Keshavan et al., 1998; Sekimoto et al., 2007), but recent studies do link reduced delta power and K-complex density in patients to cognitive deficits (Göder et al., 2006; Ramakrishnan et al., 2012) and suggest that SW abnormalities are not solely downstream of medication, since they are evident in first episode psychosis patients (Kaskie et al., 2018). These variable findings likely reflect a multitude of factors, including EEG frequency band definitions, analysis procedures, averaging power measures over multiple sleep stages and behavioral context.

To enhance sensitivity in the present study, we focused selectively on spindle and SW rich sleep stages N2 and N3, anchoring analyses to individually detected SW events. By confining spectral measures of SW amplitude and coherence to time windows surrounding these SW events, we minimized contamination by slow oscillatory noise. This approach revealed a significant reduction of frontal-occipital SW coherence in SCZ during both N2/N3 sleep. This is the first report of disrupted SW coordination in SCZ and is reminiscent of the loss of frontal-occipital SW coherence during NREM sleep in the MAM-E17 rat model (Phillips et al., 2012). Note that this reduced SW coherence is very unlikely to be secondary to changes in SW occurrence or detection, since SW densities and amplitudes were conserved in SCZ patients.

SW coherence has previously been shown to increase after learning a declarative, word-pair task (Mölle et al., 2004) and phase synchrony of SW oscillations between motor and sensory cortices has been shown to be necessary and sufficient for successful sleep-dependent consolidation of perceptual learning in rodents (Miyamoto et al., 2016). This supports a model whereby synchronized SW transitions shape distributed ensemble reactivations thought to underlie sleep-dependent memory consolidation; this coordinated activity is evidently impaired in patients with SCZ.

The UP-DOWN state transitions of cortical pyramidal neurons that underpin SW in EEG are disrupted in the MAM-E17 model, potentially reflecting a failure to temporally integrate convergent synaptic inputs (Moore et al., 2006) that may contribute to SW abnormalities in SCZ. However, SW coordination also relies on thalamocortical circuits (Crunelli and Hughes, 2010) that are compromised in SCZ (Ferrarelli and Tononi, 2011; Zhang et al., 2014). Whether the SW phenotype we report here derives from cortical, thalamocortical or corticothalamic dysfunction therefore remains an important open question.

### Disrupted SW spindle interactions in SCZ

A recent analysis of this same dataset showed that the consistency of ‘local’ SW-spindle coupling – i.e. on the same sensor – was predictive of MST memory in SCZ patients, but not in controls (Demanuele et al., 2017). By extending these analyses to cortex-wide SW-spindle interactions, we demonstrate a learning dependent increase in fronto-parietal/occipital slow wave phase to spindle amplitude coupling in healthy controls, but not in patients. Thus, while local processing during NREM sleep (Huber et al., 2004; Nir et al., 2011; Vyazovskiy et al., 2011) clearly supports aspects of sleep dependent memory consolidation (e.g. Nishida and Walker, 2007), deficits in patients with SCZ may derive from compromised top-down modulation of parietal and occipital spindle-associated activity during frontally generated SW.

The learning specific increase in fronto-occipital SW-spindle coupling is confirmed by regression analysis, showing that brain-wide SW-spindle coupling is overall the best predictor of sleep-dependent memory consolidation in healthy controls, with coupling involving parietal regions particularly prominent. A higher prediction error in patients suggests a deficit in functionally relevant SW spindle coupling.

The precise mechanisms underlying SW-phase to spindle amplitude coupling remain unclear but, given the considerable overlap of circuits involved in the generation of SW and spindles (Steriade et al., 1993; Destexhe et al., 1999; Crunelli and Hughes, 2010; Lüthi, 2014), circuit defects in schizophrenia may affect the generation and coordination of both NREM oscillations in patients (Gardner et al., 2014). Dysfunctional GABAergic transmission may be central to this SCZ phenotype; for example, the GABA receptor agonist zolpidem has been shown to enhance SW-spindle coupling and memory in healthy participants (Niknazar et al., 2015).

### Conclusions

The succession of a SW and a subsequent spindle event may not only signify local processing. We show that SW synchronization between remote cortical areas coincides with long-range SW-phase to spindle-amplitude coupling, potentially enabling coordinated information exchange between distant cortical networks and thus facilitating overnight procedural memory consolidation. SW-triggered processing can also affect the hippocampus which has been shown to be involved in some types of motor sequence learning (Schendan et al., 2003; Albouy et al., 2008; Doyon et al., 2009; Debas et al., 2010). In both rats and humans it has been shown that SW influence the timing of spindles, which are in turn synchronized to fast hippocampal ripple oscillations (Sirota et al., 2003; Mölle et al., 2006; Clemens et al., 2007, 2011; Staresina et al., 2015). This intricate synchronization of sleep oscillations might represent a mechanism for targeted information integration from the associational networks in the hippocampus during encoding to long term storage sites in the cortex (Saletin and Walker, 2012).

This precise temporal organization is impaired in patients with SCZ, hence oscillatory sleep events constitute biomarkers of circuit dysfunction (Gardner et al., 2014). In particular, using SW as markers for detailed analyses of spectral dynamics allows focus on key time windows of thalamocortical activity and function during NREM sleep, minimizing variance introduced by attention or other task variables present during wake behavior. SW and other oscillatory events during sleep provide an internal standard and a data driven approach to compare patient and healthy control data and should also frame the comparison of animal and human data in future translational studies.

## Methods

### Participants

21 schizophrenia outpatients were recruited from an urban mental health center. Diagnoses were confirmed with Structured Clinical Interviews for DSM-IV and symptoms rated according to the Positive and Negative Syndrome Scale (Kay et al., 1987; First et al., 2012). The control group of 17 healthy participants were screened to exclude a personal history of mental illness, family history of schizophrenia spectrum disorder, and psychoactive medication use. Some datasets in both CON group (baseline and learning sleep) and the SCZ group (only baseline sleep) were excluded from further analysis due to high low-frequency noise levels which affected the reliable detection of slow waves (see details of missing data and channels in the Supplementary Excel file.

The remaining patient (night 1, n=17, night 2, n=21) and control (night 1= n=15, nigth2, n=15) participants did not differ in age, sex or parental education. All participants were screened for diagnosed sleep disorders, treatment with sleep medications, a history of head injury, neurological illness and substance abuse or dependency. All participants gave written informed consent. The study was approved by the Institutional Review Boards of Massachusetts General Hospital, the Massachusetts Department of Mental Health, and Beth Israel Deaconess Medical Center.

### Sleep recordings and behavior

Participants visited the Clinical Research Center (CRC) the week before their stay to complete informed consent, demographic questionnaires, and rating scales. They also received an actigraph to wear until study completion (see Wamsley et al. 2012 for full Methods and Results).

EEG and polysomnography (PSG) data were recorded during 2 consecutive weeknights in the CRC with participants engaging in their usual activities during the intervening day. On the second night, participants were trained on the motor sequence task (MST) 1h before their usual bedtime, wired for PSG, and allowed to sleep for up to 10h. They were then tested on the MST again 1h after awakening.

### Motor Sequence Task

The MST requires pressing four numerically labelled keys in a five-element sequence (4-1-3-2-4) on a standard computer keyboard with the fingers of the left hand. The numeric sequence was displayed on a computer screen, with dots under each number indicated a keystroke. Sequences have to be entered “as quickly and accurately as possible” over 30s.

During both training and test sessions, participants had to alternate between typing and resting for 30s for a total of 12 trials, with the number of correct sequences per trial reflecting the speed and accuracy of performance. Overnight improvement was calculated as the percent increase in correct sequences from the last three training trials to the first three test trials the following morning (Fig 1C).

### Polysomnography

Different montages of five to eight channels (F3, F4, C3, Cz, C4, Pz, O1, O2) were placed according to the standard 10-20 system and EEG data digitized at 100Hz using an Embla N7000 system (Medcare Systems, Buffalo, New York). EEG was referenced to the linked mastoids for further analysis (see Supplementary Information II and Wamsley et al. 2012, for further details). Recordings were divided into 30s epochs and scored according to standard criteria (Kales et al., 1968) as WAKE, REM, N1, N2 and N3 sleep by expert scorers blind to night and diagnosis. 30 second epochs with high noise levels were discarded.

### SW and spindle detection

Sleep EEG oscillatory events were detected as described in (Phillips et al., 2012). SWs were detected from 0.25-4Hz band pass filtered EEG: the whole EEG trace (noisy epochs removed) was converted to a z-score and threshold crossings above 3.5 standard deviations (SD) from mean amplitude were detected as candidate events. These candidate events were only accepted as SW if they fell within the following parameter ranges: amplitude 50-300μV; length 0.2-3s; minimum gap between SW to be considered separate events 0.5s (Supplementary Figure S2a). All analyses presented here are based on negative singular threshold crossings with a frequency (period of oscillation) below 1.5Hz.

To detect spindles, EEG traces were band pass filtered (9–16Hz), z-scored, rectified and an envelope of the rectified signal was determined using a cubic spline fit to the maxima of the rectified signal. Candidate spindle events were detected as a threshold crossings of above 3.5 SD of the envelope, then classified as spindles given: amplitude 25-500μV; length 0.25-3s; minimum gap between spindles to be considered separate events 0.25s; start/end limit threshold 1.5 SD; oscillation frequency 12-15Hz. Differences between SW amplitudes, SW-spindle cross correlation bins and spectrogram bins were assessed using a 2-tailed Wilcoxon rank sum test with subsequent FDR correction (Benjamini and Hochberg, 1995) for multiple testing.

### EEG spectral analyses

The average spectra for Stage 2 sleep and event triggered spectral analyses were derived using multi-tapered spectra/spectrograms and coherograms using the Chronux toolbox (Mitra and Bokil, 2008, www.chronux.org). SW negative peak times were used as triggers to analyze 4s windows of EEG data (±2s around each SW). Event triggered spectrograms and coherograms were then calculated using 3 tapers, a 1s sliding data window with 50ms steps and a bandwidth of 0.1 (for SW) or 1Hz (spindle analyses) and subsequently averaged.

### Phase amplitude coupling

SW event-centered EEG windows were also used to calculate phase-amplitude coupling (PAC) using a set of previously described custom MATLAB routines (Onslow et al., 2011). We used the modulation index (MI) to quantify low frequency oscillation (0.5-5 Hz) phase to fast oscillation (5-20 Hz) amplitude coupling. The MI can quantify phase amplitude coupling within a signal or between signals from different sensors. Briefly, the modulation index is calculated as follows: both the slow frequency signal and the fast frequency signal (which can be identical if we look for PAC in the same signal, ie local PAC) are bandpass filtered at their respective frequency of interest. Next, the phase of the slow frequency signal and the amplitude of the fast frequency signal are calculated using the Hilbert transform, respectively. This results in a phase and an amplitude time series, used to create a histogram of amplitude values per phase value. If phase of the slower frequency signal had no influence on the amplitude of the faster frequency signal, we would expect a uniform distribution of amplitude values across phase bins. The modulation index thus quantifies the divergence of the amplitude distribution from a uniform distribution using a modified Kullback-Leibler distance (Canolty et al., 2006; Tort et al., 2010) where an MI of 0 indicates no phase amplitude coupling and an MI of 1 would indicate a Dirac-like distribution with all amplitude values appearing in one phase bin. We initially detected a modulation of slow wave phase (0.5-1.5 Hz) to fast spindle oscillation (12-15 Hz) amplitude using a wider frequency range (Figure 3) and subsequently ran MI calculation for SW-spindle PAC on all possible electrode pairs to quantify slow wave to fast spindle brain wide cross frequency coupling during NREM sleep.

Some analysis routines were used in combination with the Matlab© Parallel Computing Toolbox and all calculations were run on a Dell Precision Tower 7810 with two Intel^®^ Xeon^®^ CPU E5-2667 at 3.20GHz and 32GB RAM.

### Linear mixed model analysis

To analyse changes in coherence or PAC for selected electrode pairs we used general linear mixed models (Cnaan et al., 1997) to test for main and interaction effects of group and recording night. All data were entered into random intercept models with participant ID as random effects, and night and group as fixed effects using the “lme4” library (Bates and Sarkar, 2005) in the R environment (R. Core Team, 2014). Interaction terms were included if they improved the model significantly. If interaction terms were present, post-hoc multiple comparison was carried out using the “lmerTest” library (Kuznetsova et al., 2013) where p-values were calculated from F statistics of types I - III using Satterthwaite approximation of degrees of freedom.

### Data preparation and PLS regression model building

Sleep predictor variable sets to enter regression models were gathered into a wide table data format and were chosen to be average detected NREM event properties (SW and spindles) or connectivity measures such as SW coherence, SW-spindle PAC (and for comparison SW triggered spindle coherence and spindle triggered spindle coherence). We only considered measures of NREM sleep during learning night 2. The MST test results were entered as the percent change in the number of correct sequences from the last 3 trials in MST1 to the first 3 trials in MST2 (Figure 1). NREM sleep electrophysiology variables and change in MST were transformed in to z-scores (across both groups), allowing comparisons of model performance in multiples of the standard deviation for the whole population. Complete cases were then extracted for each group and modelled separately (with different levels of missingness). We compared model performances with reduced sets of variables or group sizes and the results were comparable to final models presented here.

Since the number of NREM sleep electrophysiology variables was greater than the number of individuals (p<N), partial least squares (PLS) regression (Geladi and Kowalski, 1986; Abdi, 2010; Krishnan et al., 2011) was used to reduce information in all variable sets into a smaller number of principal components (PC). PLS regression is similar to principal component regression in terms of dimensionality reduction but in PLS regression the outcome (Y) is included in the data reduction step. PLS regression works by reducing the exposure variables into PCs which have the greatest covariance with the outcome (Y). Therefore, the resulting PCs represent relevant structural information about the outcome Y (Supplementary Statistical Methods SM6).

PLS regression was carried out separately on each of the sleep exposure sets (connectivity variables) and each of the two groups (Supplementary Table S1). In order to compare prediction performance of the sleep exposure sets, it was decided *a priori* to choose 3 component models, where the explained outcome variance in the model for the EEG derived connectivity predictor X had to reach at least 80% of total variance at least in one of the groups (Wu et al., 2014). PLS models were also built with 6 components to confirm the stability of the presented results (not shown). For comparison we also present all model results for non-connectivity-based sleep exposures (NREM event properties and power measures, Supplementary Table S1) although we relaxed the 80% explained variance rule for these variable sets. The outcome (y) was taken to be the percentage overnight change in MST performance as described above. PLS regression was carried out in Matlab (Mathworks, Natick MA) using the built-in function *plsregress*. The residual sum of squares (RESSS) and R^2^ were used to compare models between groups and exposure sets (Abdi, 2010).

We used permutation tests to assess the whether the difference in RESS values between groups is likely to significant. We computed 10,000 permutations of group membership (CON, SCZ) with fixed group sizes to estimate the null distribution, i.e. no difference in RESS between randomly composed groups (dRESS=0). We then computed a PLSR model for each of the permutations and each variable set to compare the sample difference in RESS with the null distribution. Two-sided p values were calculated as the number of absolute dRESS values that are bigger than the sample value divided by total number of permutations. Given we tested 15 sleep variable sets, we applied a Bonferroni threshold of 0.005/15= 0.0033 as significance threshold.

## Supporting information

Supplemental Information - Supplemental Results, Tables, Statistics

Supplemental Information 2 - Missing data

## Acknowledgments

We indebted to Dara Manoach (Department of Psychiatry, Massachusetts General Hospital) and Robert Stickgold (Department of Psychiatry, Beth Israel Deaconess Medical Center) for generous sharing of data recorded in their labs and for comments on early drafts of the manuscript. UB and MWJ thank the Medical Research Council (UK) for support (Fellowship G1002064). UB thanks Lilly UK for support in form of Lilly Innovation Fellowship Award.

## Author contributions

UB coded and ran all analyses, generated figures and drafted the manuscript; AS designed and ran statistical tests with UB; CD discussed earlier stages of the project and commented on the manuscript; EW recorded all the data and commented on the manuscript; HM advised on the approach and commented on the manuscript; MWJ conceived the study alongside UB and Dara Manoach and co-wrote the manuscript.

## Competing financial interests

UB received full time salary through Lilly UK, as part of a Lilly Innovation Fellowship Award.

**Materials & Correspondence**

## Abbreviations

SW: slow wave
SCZ: schizophrenia
MST: motor sequence task
PSG: polysomnography

