## Supplemental Information - Supplemental Results, Tables, Statistics for "Neurophysiological predictors of memory impairment in schizophrenia: decoupled slow-wave dynamics during non-REM sleep"

### Supplementary Results

#### SR1 - Overall spectral properties of NREM sleep

We characterized the overall spectral properties of NREM sleep epochs (all N2 and N3 epochs) in both groups during both recording nights using multi-taper spectra. Figure S1a and b show the average spatial distribution of SW and spindle power during baseline and learning nights, respectively in healthy control and patients.

Figure S1c shows the average spectra at each electrode averaged over both nights for healthy controls and patients. Individual frequency bins were tested with a Wilcoxon rank sum test and all p values were corrected according to the Benjamini-Hochberg procedure (implemented in Matlab mafdr.m. referred to as p_BH_ below) to account for false discovery rates. Surviving bins were marked with a green tick. Notice the significant differences in spindle power and lower frequencies in patients.

#### SR2 - Slow wave detection and slow wave properties during NREM sleep

We detected slow wave events using a custom Matlab® routines. Figure S2a shows an example slow wave detection. Slow wave events were detected using the criteria described in the main text methods. Figure S2b shows wave triggered averages for all electrodes for both healthy volunteers (CON) and patients (SCZ).

Individual time bins were tested with Wilcoxon ranksum text and p values were corrected according to the Benjamini-Hochberg procedure. Significant bins (p_BH_ <0.05) were marked with a green tick.

Figure S2c shows histograms of slow wave properties that are amplitude, intrinsic frequency and length in time (duration) and slope normalized to the total number of events for both groups and both recording nights. Testing differences in distributions with a Kolmogorov Smirnoff test did not reveal any significant differences in distributions (p<0.0125).

#### SR3 - Slow wave densities

Figure S3 shows slow wave densities at all sensor locations. There is no difference in slow wave density between healthy volunteers and patients on either recording night (p_BH_ > 0.05, 2-sided Wilcoxon rank sum test, FDR corrected).

#### SR4 - Spindle detection and spindle properties during NREM sleep

We detected spindle events using a custom Matlab® routine (Phillips et al., 2012). Figure S4a shows an example spindle detection, spindle events were detected using the criteria described in the main text methods. Figure S4b shows wave triggered averages for all electrodes for both healthy volunteers (CON) and patients (SCZ). Significantly different bins were marked with a green tick (p_BH_ <0.05, 2-sided Wilcoxon rank sum test, FDR corrected).

Figure S4c shows histograms of spindle properties that are amplitude, intrinsic frequency and length in time (duration) normalized to the total number of events for both groups and both recording nights. Testing differences in distributions with a Kolmogorov Smirnoff test did not reveal any significant differences in distributions (p<0.0125).

#### SR5 - Spindle densities

Figure S5 shows spindle densities at all sensor locations. There are significant differences in spindle density between healthy controls and patients (p_BH_ <0.05, 2-sided Wilcoxon rank sum test, FDR corrected) but not between recording nights.

#### SR6 - Slow wave spindle interactions on the same sensor I

To examine distributed coupling of SW and spindle we first calculated histograms of spindle time triggered on SW trough times. Figure S6a shows SW trough time triggered histograms of spindle start times at each sensor. Overall correlation between SW and spindles is reduced in patients compared healthy controls although significant differences in time bins only appear at Cz (marked by green ticks, pBH<0.05, 2-sided Wilcoxon rank sum test, FDR corrected

Next, we calculated multi-taper spectrograms using negative peak of any detected SW (wave triggered average overlaid in white) as reference. The resulting spectrograms were averaged across nights for each group (Figure S6b). In the CON group (upper two rows) there is clear modulation of spindle power (12-15 Hz) relative to SW. The trough of the SW (DOWN state) is flanked by two high spindle power periods, before and after (peak, UP state). Highest local SW associated spindle power is observed at central (Cz) and parietal (Pz) recording sites. Figure S6c shows the average spindle power calculated from SW triggered multi-tapered spectrograms (-2-2sec, 12-15Hz). Stars indicate a significant difference between CON and SCZ group (p_BH_ < 0.05, 2-sided Wilcoxon rank sum text with FDR).

#### SR7 - Slow wave spindle interactions on the same sensor II: SW-spindle PAC at Cz

We calculated the modulation index as described in the main methods from ±2s windows surrounding SW negative peak times of SWs recorded at Cz. Notice the strong SW (0.5-1 Hz) phase to fast spindle (12-15 Hz) amplitude modulation present in healthy controls on both recording nights, but less so in patients. Indeed linear mixed model analysis of average modulation index (0.5-1.5 Hz, 12-15Hz) reveled a significant group effect (F_(1, 34.137)_= 18.59, p=0.00013) but not for night (F_(1,29.446)_= 0.099 p=0.75, see also Supplementary M4). No interaction effect was detected as evident in the interaction plot (Figure S7b).

#### SR8 - Slow wave triggered spindle coherence

Coordination of SW-nested spindles across different electrode positions was quantified by calculating coherence in ±2s windows surrounding SW negative peak times (Figure S8).

Figure S8a shows example coherogram for the pair Cz O1 for both groups on baseline and learning nights. Spindle coherence appears to peak near the peaks of detected slow waves (wave triggered average overlaid in black), in accordance with previously shown spectrograms that show SW-time locked increases in spindle amplitude. SCZ patients showed attenuated SW-modulated centro-occipital spindle coherence during both nights.

Indeed, linear mixed model analysis of average Cz-O1 SW-triggered spindle coherence values (-2-2 s, 12-15 Hz) revealed significant effects of group (F_(1, 34.705)_=8.04, p=0.0076), and a trend for night (F_(1,30.338)_=3.24 p=0.082, Figure S8b) but no interaction term.

SW-modulated spindle coherence matrices (12-15Hz, Figure S8c) showed a similar pattern to SW coherence. Coherence with parietal regions was high for both occipital and front-central regions, suggesting Pz acts as a link between fronto-central networks and occipital networks. SW-associated spindle coherence was lower in SCZ patients than controls during the baseline night across many electrodes, including front/central-occipital and parietal-occipital electrode pairs (Figure S8c, Δ_[CON – SCZ base]_, bottom, left). Learning induced increases in central to occipital coupling via SW-associated spindle coherence in healthy controls (Figure S8c, Δ_[CON: learn – base]_, top, right). In contrast, learning related changes in SCZ are less pronounced compared to healthy controls (Figure S8c, Δ_[SCZ: learn – base]_, right, middle).

#### SR9 - Spindle triggered spindle coherence

Coordination between detected spindle events across different recording locations was quantified by calculating coherence in ±2s windows surrounding spindle maximum peak times (Figure S9). In the Control group, there was strong spindle coherence between central and occipital regions (Figure S9a, top right plot). In contrast, SCZ patients showed attenuated centro-occipital spindle coherence during both nights.

Linear mixed model analysis of average Cz-O1 spindle coherence values (-2-2 s, 12-15Hz) revealed significant effects of group (F_(1,34.845)_= 9.52, p=0.004), but not for night (F_(1,29.927)_ =2.84 p=0.10, Satterwaithe approximation of degrees of freedom) Figure S9b shows the least-squares estimated group averages on both nights in an interaction plot.

Spindle coherence matrices (Figure 9c) showed similar patterns to those evident for SW triggered spindle coherence but with higher overall coherence values. Overall spindle coherence was lower in SCZ patients compared controls during both baseline and learning nights across mainly fronto/central-ocipital electrode pairs (Figure S9c, Δ_[CON – SCZ base]_, bottom). Learning related changes in overall spindle coherence in both groups were small (Figure S9c, Δ_[CON: learn – base]_, Δ_[SCZ: learn – base]_, right). In summary, overall spindle coherence is impaired in SCZ with fewer learning dependent changes.

#### SR10 - PLS and linear regression of NREM sleep features

Table S1 shows the final prediction error (RESS), R squared for each PLSR model and F and p-value for the subsequent stepwise linear model fit for each variable set. A variable set’s predictive properties are quantified by their RESS value. Due some datasets not meeting inclusion criteria at individual electrodes (mostly occipital electrodes) complete cases N numbers for the different models varied. All models with local SW and spindle properties had the same number of participants entered: SW density, SW amplitude SW frequency SW length SW slope spindle density spindle amplitude, spindle frequency, spindle length: N_CON_ = 10, N_SCZ_ =14. For all other variables N numbers were: spindle triggered spindle coherence: N_CON_ = 11, N_SCZ_ =10, SW triggered spindle coherence: N_CON_ = 8, N_SCZ_ =11, SW coherence: N_CON_ = 8, N_SCZ_ =14, SW spindle power N_CON_ = 8, N_SCZ_ =13, SW spindle PAC: N_CON_ = 7, N_SCZ_ =13.

The top half of the table shows models for properties of detected events. These include SW density during NREM (events/min), SW amplitude (μV), SW frequency (1/ peak to trough time difference), SW length (peak to peak time difference) SW maximum slope (μV/s) and spindle density during NREM (events/min), spindle amplitude (μV), spindle frequency (1/ peak to trough time difference), spindle length (peak to peak time difference). Here, SW properties are overall better predictors of successful sleep dependent memory consolidation compared to spindle properties (in both healthy controls and patients). Some SW features are more predictive in controls (SW length and slope) whereas SW density may be more predictive in patients. Spindle features are less predictive of sleep dependent memory consolidation, but spindle density is slightly more predictive in healthy control but not in patients although not of the subsequent linear models reach significance.

Of the connectivity measures SW spindle PAC is the best predictor of sleep dependent memory consolidation (see main text) but all SW related connectivity measures reach similarly low RESS values in controls but not in patients. Only spindle triggered spindle coherence appears to be a better predictor in patients, suggesting that spindle coherence is more relevant for memory processing in patients compared to controls. Overall, SW coherence and measures of SW spindle coordination are better predictors of memory consolidation in controls compared to patients implying that functional deficits in SW and SW-spindle coordination underlie deficits in sleep dependent memory consolidation in patients.

### Supplementary Figures

#### Figure S1: Overall spectral properties of NREM sleep.

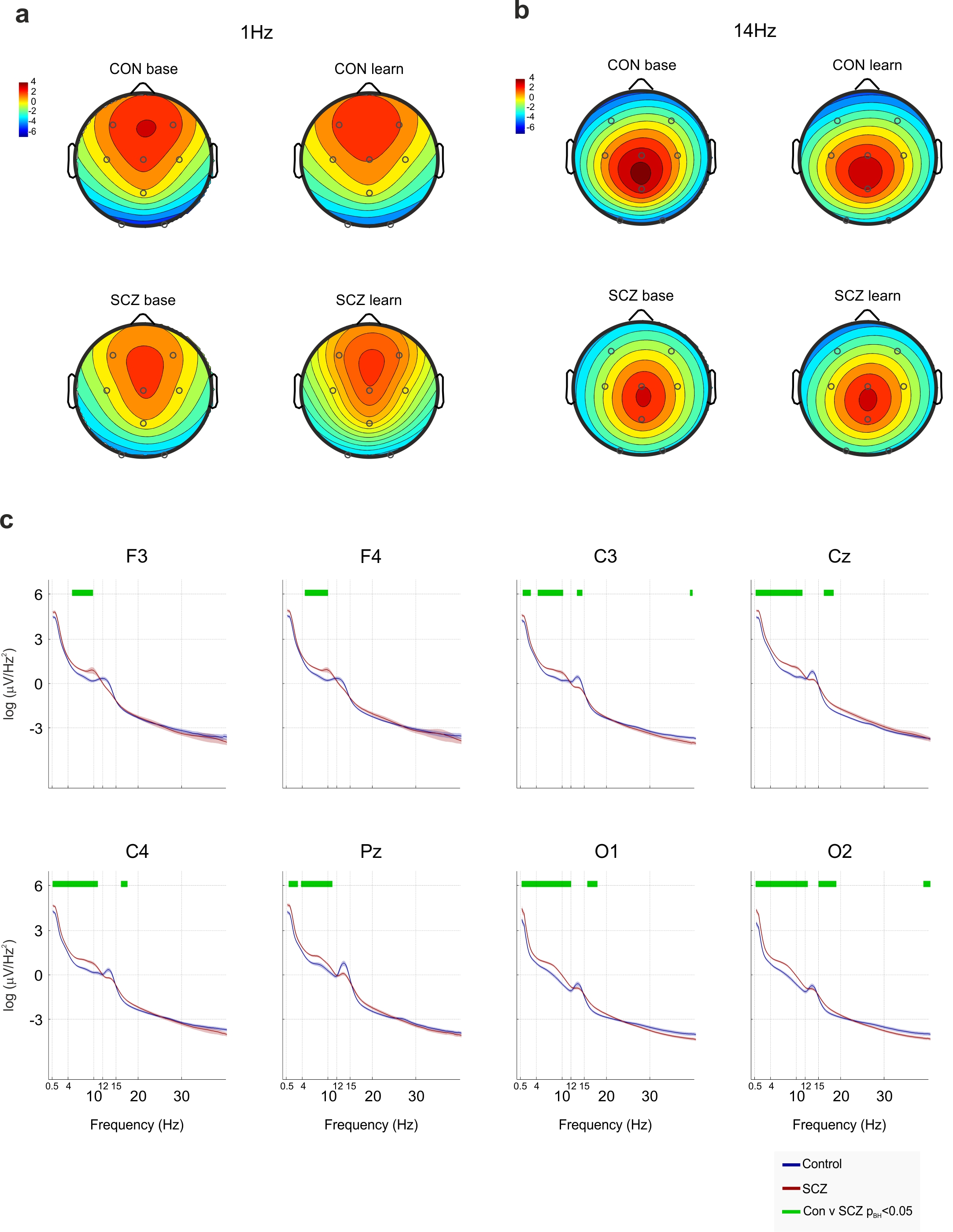

a) Average peak SW power

Sensor-wide distribution of peak SW power (1Hz) for both groups (SCZ, CON) and both recording nights (baseline and learning) averaged from all N2 sleep epochs.

b) Average peak spindle power

Sensor wide distribution of peak spindle power (14Hz) for both groups (SCZ, CON) and both recording nights (baseline and learning) averaged from all N2 sleep epochs.

c) Average multi-tapered power spectra at all electrode positions

Averaged multi-tapered spectra at each sensor for both groups (SCZ, CON, averaged for both recording nights) averaged from all NREM sleep epochs. Significantly different frequency ranges between CON and SCZ are marked by green ticks above the spectra (CON vs SCZ, p_BH_< 0.05, 2-tailed Wilcoxon rank sum test with FDR correction).

#### Figure S2: Slow wave detection and slow wave properties during NREM sleep

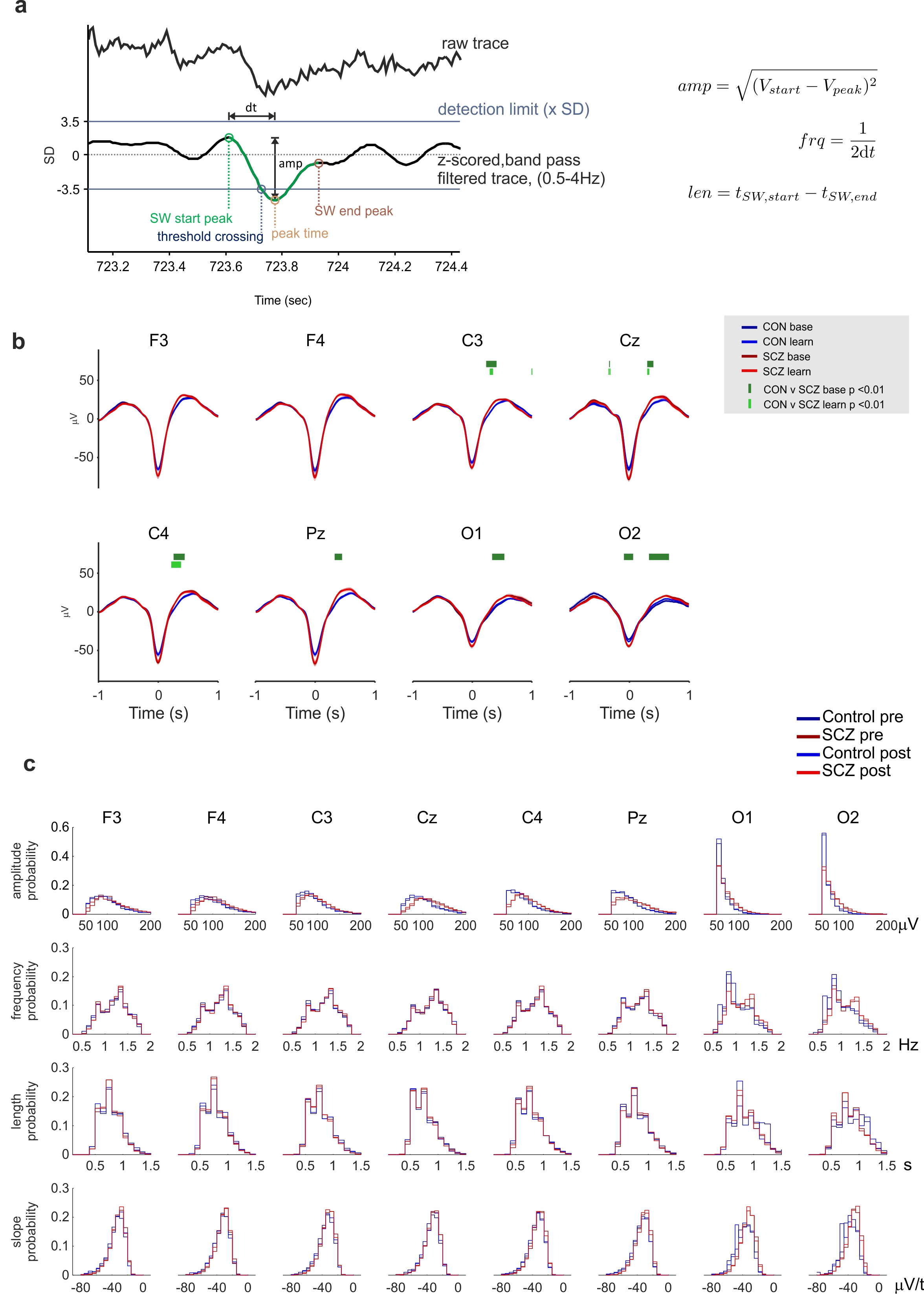

a) Slow wave activity detection algorithm.

Individual slow waves were detected based on threshold crossings of a z-scored, band pass filtered (0.5-4Hz) EEG (Phillips et al., 2012). Properties of events (amplitude, frequency, length) were calculated as described in the plot (formulae on the right).

b) Mean z-scored wave triggered averages of slow waves

SWs with frequency between 0.5 and 1.5 Hz, using negative peak time as trigger. Significantly different time bins between CON and SCZ are indicated by green ticks (dark green, higher, CON vs SCZ base, p< 0.05, 2-tailed Wilcoxon rank sum test; dark green, lower, CON vs SCZ learn, p< 0.05, 2-tailed Wilcoxon rank sum test with FDR correction).

c) Average histograms of slow wave properties

Histograms of SW properties were calculated per participant and averaged per sensor for both groups (CON and SCZ) and both nights (baseline and learning): top SW amplitude (μV) distribution, middle frequency distribution (Hz), bottom length distributions (s).

#### Figure S3: Slow wave densities

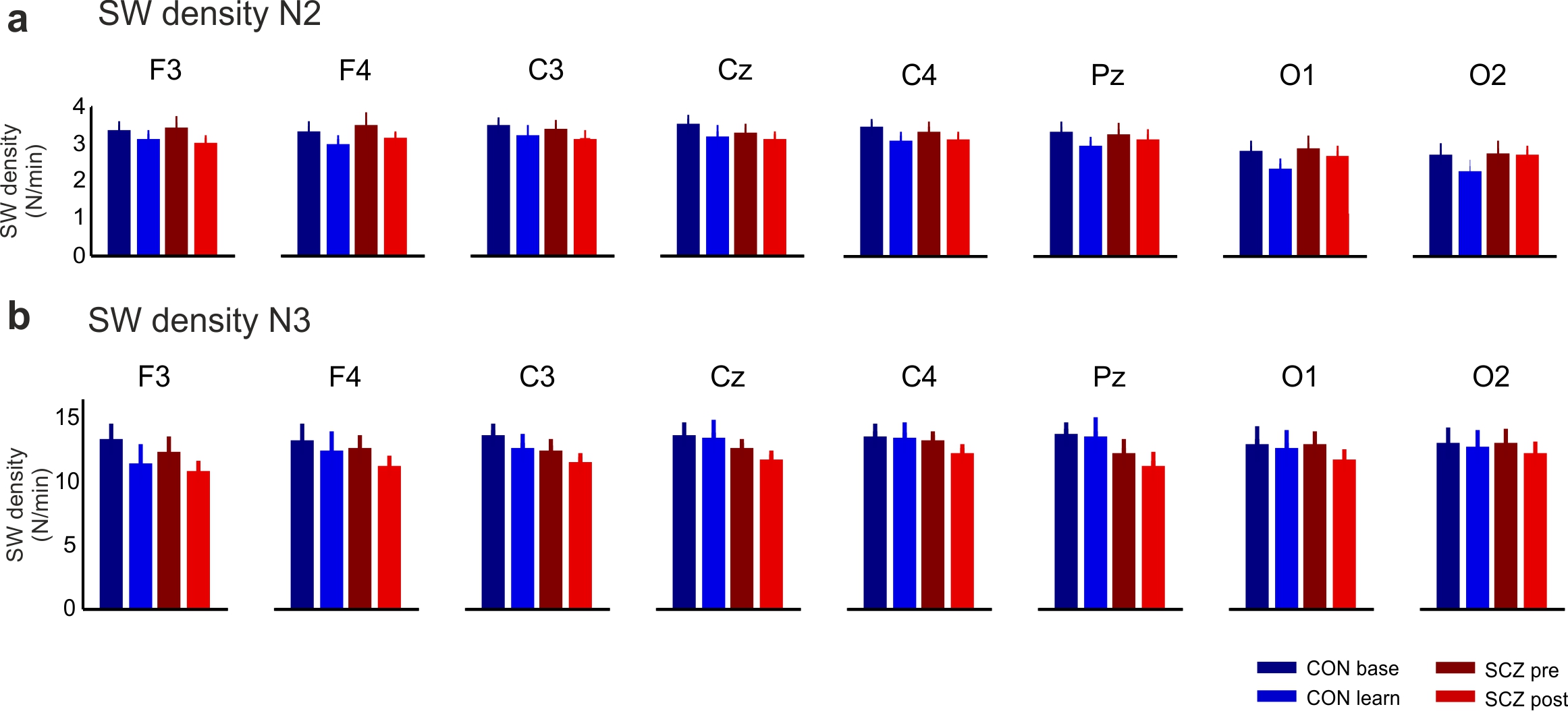

a) Average cumulative SW amplitude distributions

Histograms of SW amplitudes were calculated per participant and averaged per sensor for both groups (CON and SCZ) and both nights (baseline and learning) and plotted as cumulative sums. A two-sided Kolmogorov-Smirnov test was used to test whether the resulting distribution functions are likely to come from the same distribution or not (* p<0.05, two-sided KS test).

b) Slow wave densities per N2 and N3

SW densities (N_SW_/min) at all recording sites for both CON and SCZ during baseline and learning sleep. No significant differences were observed (alpha = 0.05, 2-sided Wilcoxon rank sum text with FDR).

#### Figure S4: Spindle detection and properties

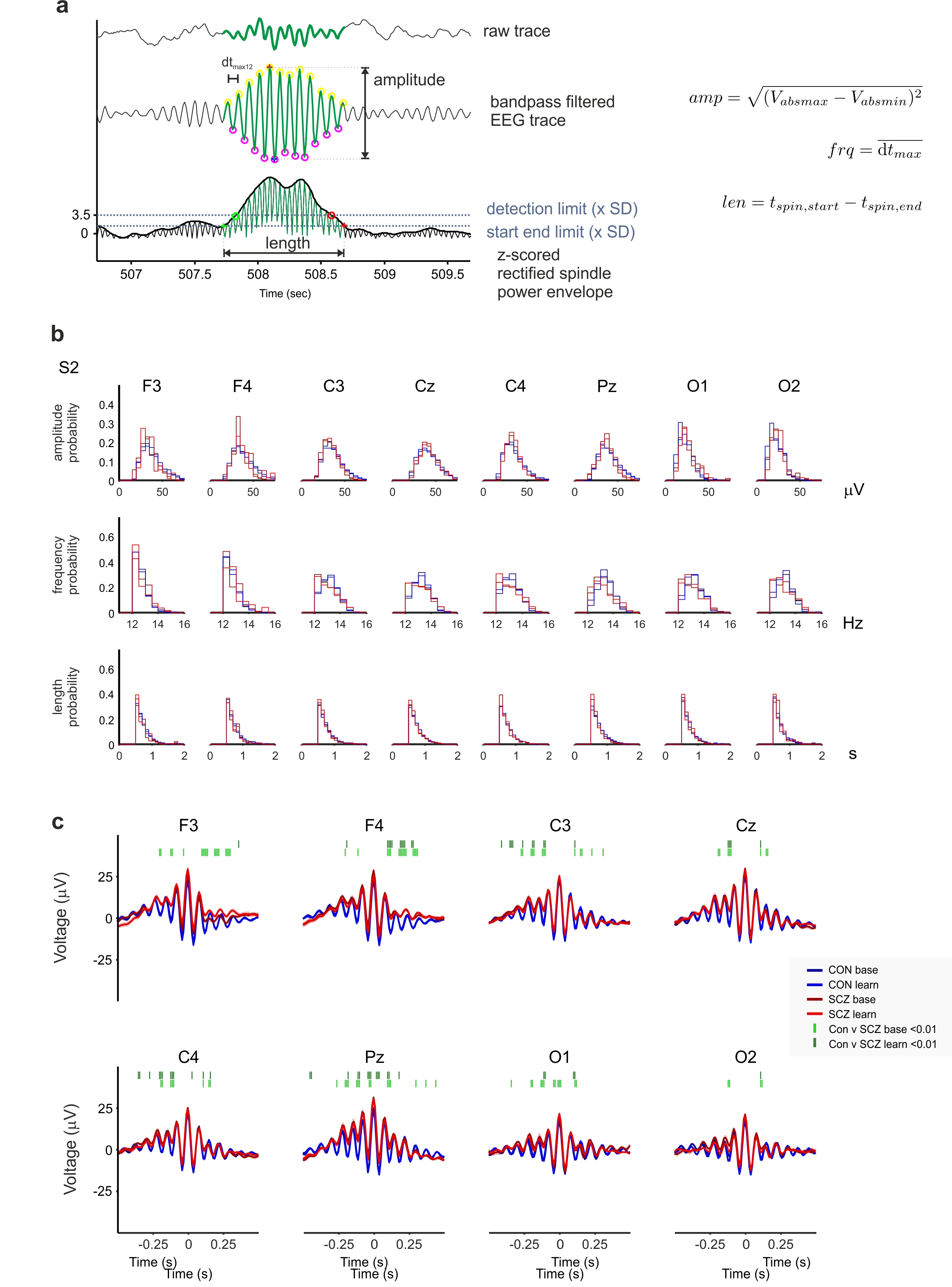

a) Automated spindle detection

Individual spindle events were detected based on threshold crossings of a spindle power envelope. The raw EEG trace was band pass filtered (9-16 Hz), rectified, and envelope of the signal was defined by cubic spline interpolation between the maxima of the rectified signal (Phillips et al., 2012). Properties of events (amplitude, frequency, length) were calculated as described in the plot (formulas on the right).

b) Average histograms of spindle event properties

Histograms of spindle event properties were calculated per participant and averaged per sensor for both groups (CON and SCZ) and both nights (baseline and learning): top, SW amplitude (uV) distribution, middle, frequency distribution (Hz), bottom, length distributions (s).

c) Mean z-scored wave triggered averages of spindles

Spindle wave triggered averages with frequency between 12 and 15 Hz, using maximum peak time as trigger. Significantly different time bins between CON and SCZ are indicated by green ticks (dark green, higher, CON vs SCZ baseline, dark green, lower, CON vs SCZ learning, p< 0.05, 2-tailed Wilcoxon rank sum test with FDR correction).

#### Figure S5: Spindle densities

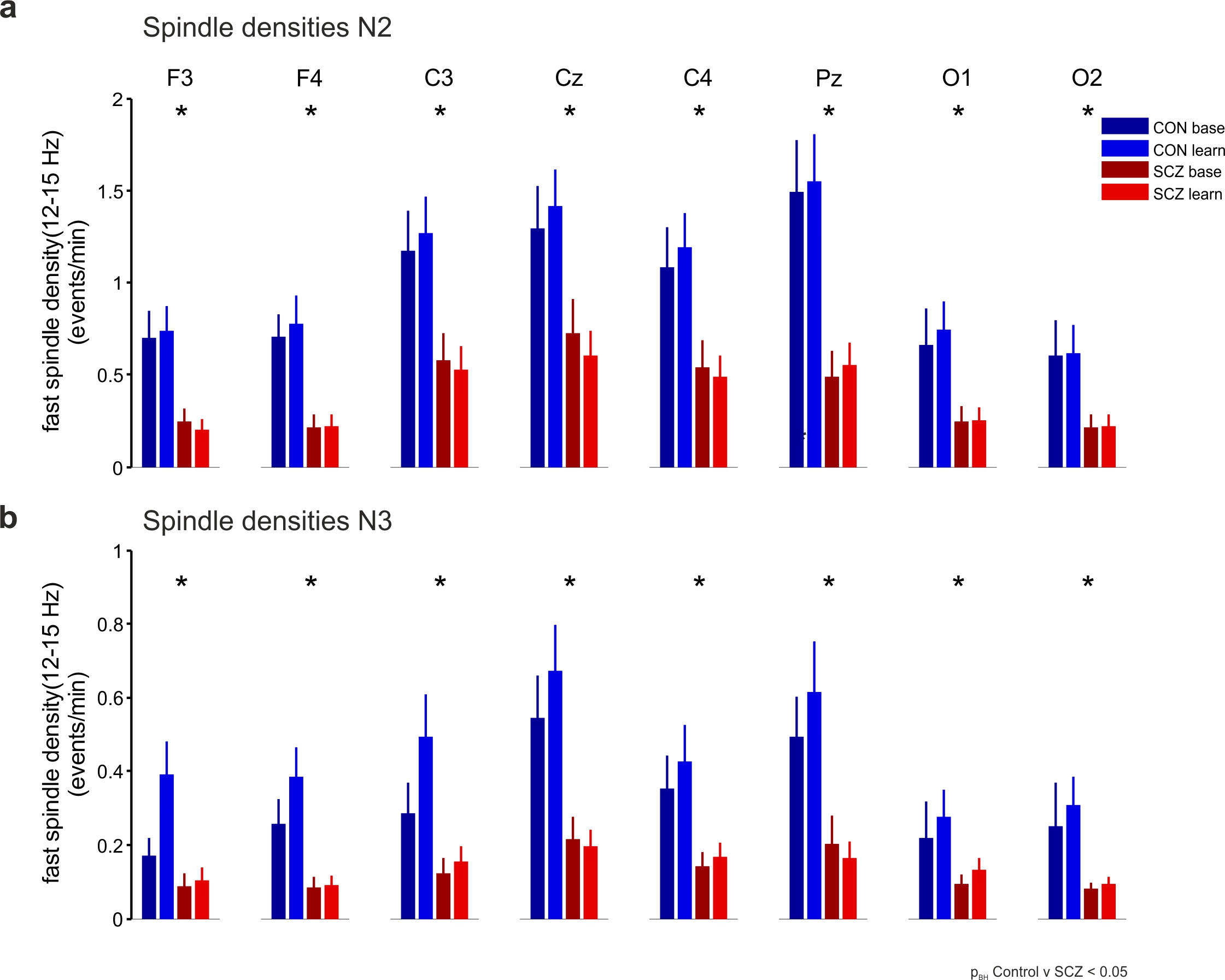

a) N2 sleep spindle densities

Spindle densities (12-15 Hz) during N2 sleep at all recording sites for both CON and SCZ during baseline and learning sleep. Stars indicate significant differences (p_BH_ < 0.05, 2-sided Wilcoxon rank sum text with FDR).

b) N3 sleep spindle densities

Spindle densities (12-15 Hz) during N3 sleep at all recording sites for both CON and SCZ during baseline and learning sleep. Stars indicate significant differences (p_BH_ < 0.05, 2-sided Wilcoxon rank sum text with FDR).

#### Figure S6: SW spindle interactions on the same sensor I

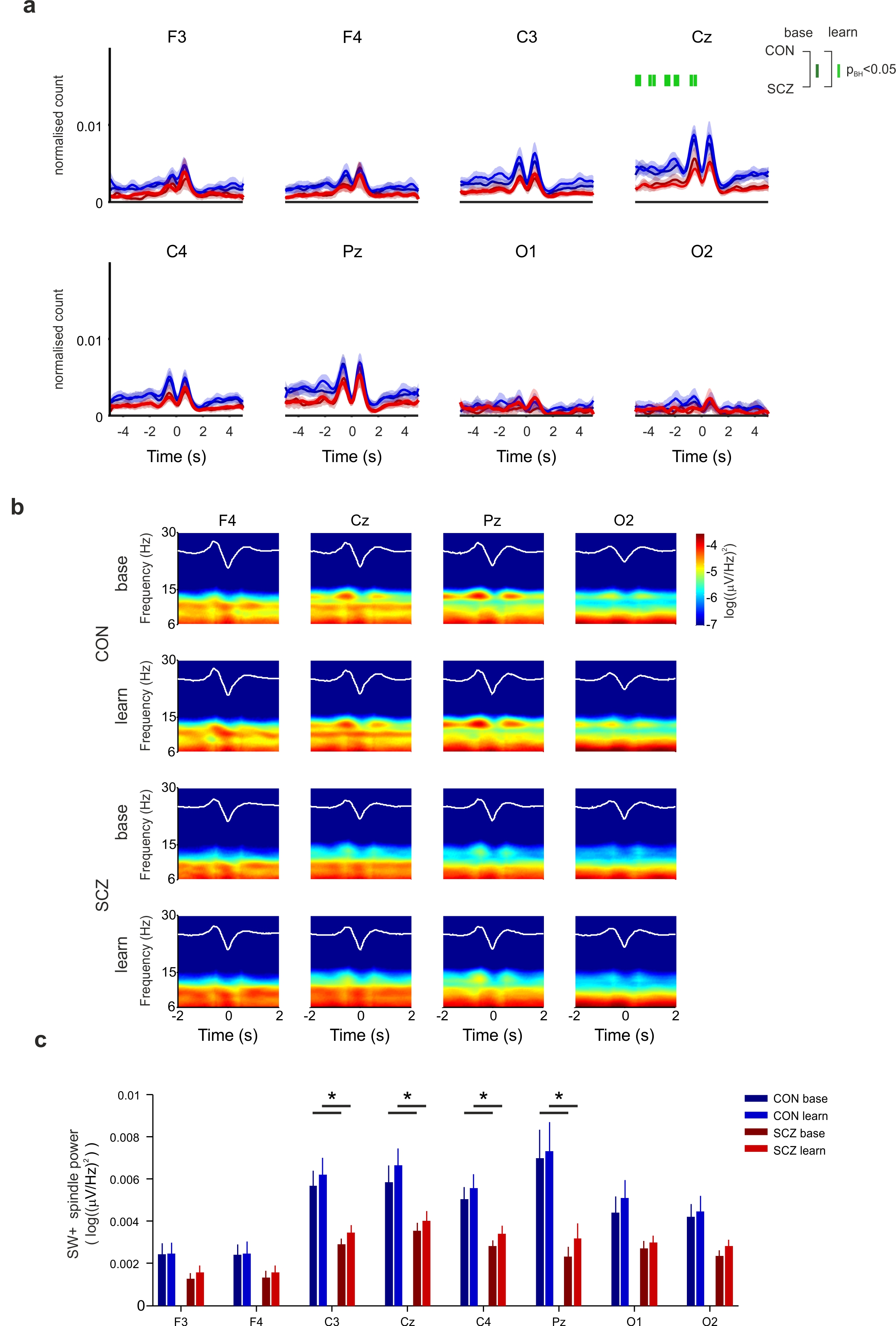

a) SW-spindle event times cross correlations

SW peak times were used to calculate cross correlations with spindle event occurrences (spindle start times). Significantly different time bins between CON and SCZ are indicated by green ticks (dark green, CON vs SCZ base, p< 0.05, 2-tailed Wilcoxon rank sum test; light green, CON vs SCZ learn, p< 0.05, 2-tailed Wilcoxon rank sum test, FDR corrected). Significant differences occur only during the learning at Cz.

b) SW triggered spectrograms

The negative peak of any detected SW (wave triggered average overlaid in white) was used as reference point to calculate sliding window multi-tapered spectrograms during a surrounding ±2s window of data (Each SW constitutes a “single trial”). The resulting spectrograms were averaged across participant groups (CON, SCZ) for each night (baseline and learning). In the control group (upper two rows) there is clear modulation of spindle power (12-15 Hz) relative to SW. The negativity of the SW (DOWN state) is flanked by two high spindle power periods, before and after (UP state). Highest SW associated spindle power can be observed at central (Cz) and parietal (Pz) recording sites but less so at frontal and occipital sites.

c) Average slow wave associated spindle power

The average spindle power was calculated from SW triggered multi-tapered spectrograms (-2-2sec, 12-15Hz). Stars indicate a significant difference between CON and SCZ group (p_BH_ < 0.05, 2-sided Wilcoxon rank sum text with FDR).

#### Figure S7: SW spindle coherence

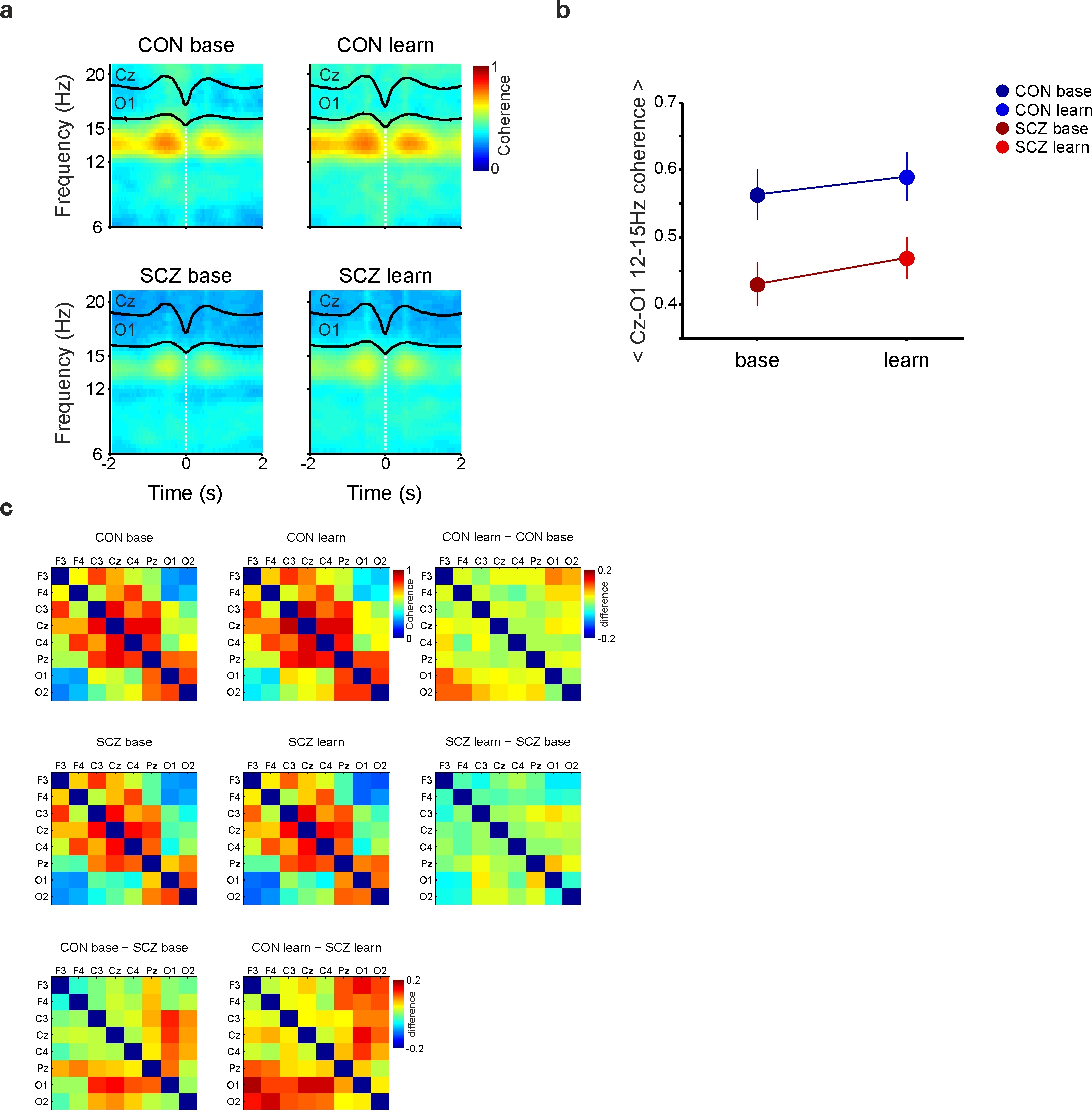

a) SW-triggered spindle coherograms.

SWs were detected at Cz, and a ±2s window around the negative peak times of SWs was used to compute a sliding window multi-taper coherogram between Cz and O1. The resulting coherograms were averaged per group and recording night. Wave triggered averages of SWs at the trigger channel (Cz, black line, top) and target channel (O1, black line, bottom) are overlaid. Spindle oscillations appear to be most coherent near positivity of a SW (UP state). Control subjects show robust spindle coherence near SWs during baseline sleep and a slight increase in spindle coherence during learning sleep although these effects are not significant (see interaction plot below). In the SCZ group, there is a distinct reduction of SW associated spindle coherence during both nights compared to CON group and a slight learning dependent increase although this does not reach significance either (see interaction plot below).

b) Interaction plot of mean SW associated spindle coherence

The mean SW-modulated coherence values from a) were fed into a linear mixed model analysis. The interaction plot shows least squares estimates of average SW-modulated coherence for both recording nights. Average coherence values significantly differ between groups but not between nights (Supplementary Statistics, SM5).

c) Brain wide SW associated spindle coherence (12-15 Hz).

Coherence matrices display SW associated spindle coherence (12-15 Hz) between all sensor pairs. Mean coherence values (12-15 Hz, ±2 s) were plotted against electrode position to create a coherence matrix. Average coherence matrices were calculated for baseline (left) and learning night (right) for each participant group (CON, top; SCZ, bottom), respectively. Differences in SW associated spindle coherence between baseline and learning night for both groups were added to the right and below the respective matrices of mean values. Note the strong differences between CON and SCZ group on both recording nights (bottom matrices) and the increase in differences during the learning night, particularly between frontal/central-occipital electrode pairs.

#### Figure S8: Spindle triggered spindle coherence

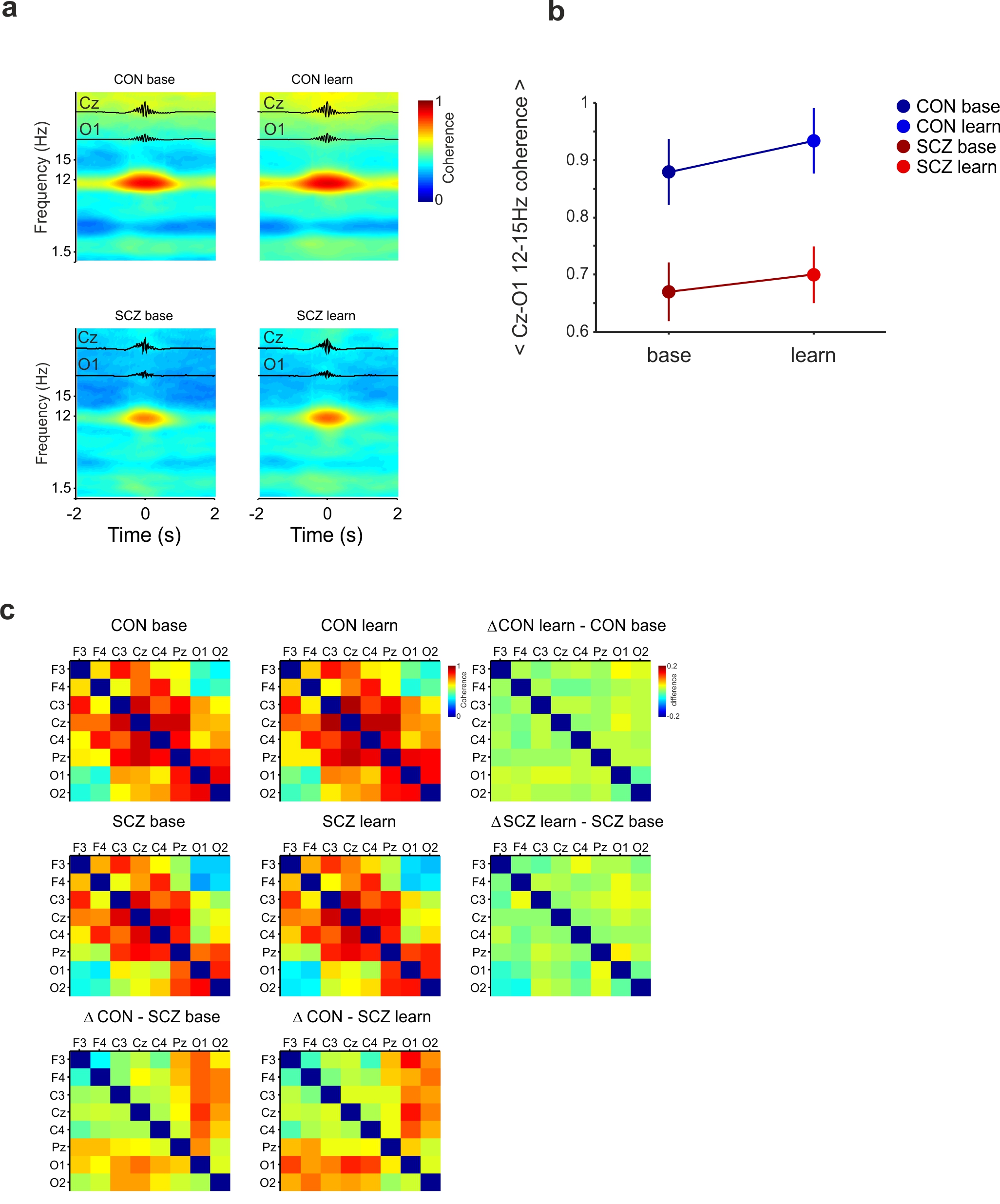

a) Spindle-triggered coherence

Spindles were detected at Cz, and a ±2s window around the negative peak times of spindles was used to compute a sliding window multi-taper coherogram between Cz and O1. Wave triggered averages of spindles at the trigger channel (Cz, black line, top) and target channel (O1, black line, bottom) are overlaid (aligned to absolute maximum amplitude). Spindle oscillations appear to be most coherent near the absolute maximum amplitude. In the SCZ group, there is little change between recording nights but distinct reduction of spindle coherence during either night compared to CON group learning dependent increase although this does not reach significance either (see interaction plot and coherence matrices below).

b) Interaction plot of mean Cz-O1 spindle coherence

The mean spindle coherence values (12-15 Hz, ±1 s) displayed in a) were fed in to linear mixed effects model. The interaction plot demonstrates the lack of a significant interaction effect between recording night and participant group in the final model. Average coherence values differ between groups but not between recording nights (see Supplementary Statistical Methods, SM6).

c) Brain wide spindle coherence (12-15 Hz)

Mean coherence values (12-15 Hz, ±1 s) were plotted against electrode position to create a coherence matrix. Average coherence matrices were calculated for baseline (left) and learning night (right) for each participant group (CON, top; SCZ, bottom), respectively. Differences in spindle coherence between baseline and learning night for both groups were added to the right and below the respective matrices of mean values. Note the group differences between CON and SCZ group on both recording nights (bottom matrices) but little change between recording nights in both groups.

### Supplementary Tables

#### Table S1: PLSR model results for all variable sets

|  | **PLSR statistics** | | | | | | |
| --- | --- | --- | --- | --- | --- | --- | --- |
|  | **SCZ** | | | **SCZ** | |  | |
| **EEG variable** | **R^2^** | **RESS** | **R^2^** | | **RESS** | | **p** |
| SW density | 0.43 | 5.13 | 0.57 | | 5.54 | | 0.88 |
| SW amplitude | 0.30 | 6.28 | 0.48 | | 6.76 | | 0.84 |
| SW frequency | 0.63 | 3.35 | 0.42 | | 7.52 | | 0.049 |
| SW length | 0.74 | 2.34 | 0.46 | | 6.98 | | 0.019 |
| SW slope | 0.88 | 1.06 | 0.31 | | 8.94 | | **1.0e-4 *** |
| spindle density | 0.72 | 2.48 | 0.29 | | 9.26 | | **6.0e-4 *** |
| spindle amplitude | 0.33 | 6.05 | 0.30 | | 9.12 | | 0.15 |
| spindle frequency | 0.43 | 5.09 | 0.39 | | 7.87 | | 0.35 |
| spindle length | 0.39 | 5.48 | 0.44 | | 7.29 | | 0.49 |
| SW coherence | 0.86 | 0.96 | 0.63 | | 4.75 | | **2.0 e-4 *** |
| spindle triggered  spindle coherence | 0.65 | 3.46 | 0.85 | | 1.37 | | 0.22 |
| SW triggered  spindle coherence | 0.88 | 0.83 | 0.85 | | 1.84 | | 0.18 |
| SW local spindle power | 0.88 | 0.81 | 0.55 | | 5.42 | | **5.0 e-4*** |
| SW remote spindle power | 0.75 | 2.23 | 0.44 | | 7.25 | | **6.0 e-4*** |
| SW spindle PAC | 0.94 | 0.34 | 0.75 | | 2.99 | | **5.0 e-4 *** |

* p value from permutation test, (n=10000), significant difference in RESS between CON and SCZ,

at Bonferroni corrected alpha = 0.0033

### Supplementary Statistical Methods

##### SM1 LMM for slow wave triggered slow coherence (Figure 2)

```
Model comparison fixed effects only v interaction:

     0: dcoh ~ night + group + (1 | id)
     1: dcoh ~ night * group + (1 | id)

       Df     AIC     BIC logLik deviance  Chisq Chi Df Pr(>Chisq)
  0     5 -41.837 -32.804 25.919  -51.837
..1     6 -44.681 -33.841 28.340  -56.681 4.8436      1    0.02775 *
```

```
Linear mixed model fit by REML
t-tests use  Satterthwaite approximations to degrees of freedom ['lmerMod']
Formula: dcoh ~ night * group + (1 | id)

Analysis of Variance Table of type III  with  Satterthwaite
approximation for degrees of freedom
              Sum Sq  Mean Sq NumDF  DenDF F.value Pr(>F)
night       0.013127 0.013127     1 20.630  1.5576 0.2260
group       0.012115 0.012115     1 23.985  1.4375 0.2423
night:group 0.042509 0.042509     1 20.630  5.0441 0.0358 *
---
Signif. codes:  0 ‘***’ 0.001 ‘**’ 0.01 ‘*’ 0.05 ‘.’ 0.1 ‘ ’ 1
```

```
Differences of LSMEANS:

                            Estimate Standard Error   DF t-value Lower CI Upper CI p-value
night:group  1 HC -  2 HC       -0.1         0.0426 20.6   -2.35  -0.1885  -0.0113    0.03 *
night:group  1 HC -  1 SCZ       0.0         0.0649 35.6    0.03  -0.1296   0.1337    0.97
night:group  1 HC -  2 SCZ       0.0         0.0616 32.4    0.49  -0.0951   0.1559    0.63
night:group  2 HC -  1 SCZ       0.1         0.0649 35.6    1.57  -0.0297   0.2336    0.12
night:group  2 HC -  2 SCZ       0.1         0.0616 32.4    2.11   0.0048   0.2557    0.04 *
night:group  1 SCZ -  2 SCZ      0.0         0.0400 21.3    0.71  -0.0549   0.1115    0.49
```

##### SM2 LMM for F3 SW triggered F3-O1 PAC (Figure3)

```
Model comparison fixed effects only v interaction:
object: swspinpac ~ night + group + (1 | ID)
   ..1: swspinpac ~ night * group + (1 | ID)

       Df     AIC     BIC logLik deviance  Chisq Chi Df Pr(>Chisq)
object  5 -123.28 -114.25 66.640  -133.28
..1     6 -125.69 -114.85 68.846  -137.69 4.4129      1    0.03567 *
```

**Linear mixed model fit by REML**

t-tests use Satterthwaite approximations to degrees of freedom ['lmerMod']

Formula: swspinpac ~ night * group + (1 | ID)

```
Analysis of Variance Table of type III  with  Satterthwaite
approximation for degrees of freedom

               Sum Sq   Mean Sq NumDF  DenDF F.value  Pr(>F)
night       0.0058587 0.0058587     1 18.626  5.3010 0.03304 *
group       0.0063891 0.0063891     1 22.540  5.7809 0.02483 *
night:group 0.0052141 0.0052141     1 18.626  4.7178 0.04298 *
```

```
Differences of LSMEANS:
                            Estimate Standard Error   DF t-value Lower CI Upper CI p-value
night:group  1 HC -  2 HC        0.0         0.0155 18.6   -3.02  -0.0790  -0.0142   0.007 **
night:group  1 HC -  1 SCZ       0.0         0.0275 30.9    1.37  -0.0184   0.0936   0.181
night:group  1 HC -  2 SCZ       0.0         0.0268 28.9    1.35  -0.0185   0.0910   0.186
night:group  2 HC -  1 SCZ       0.1         0.0275 30.9    3.07   0.0282   0.1403   0.004 **
night:group  2 HC -  2 SCZ       0.1         0.0268 28.9    3.09   0.0281   0.1376   0.004 **
night:group  1 SCZ -  2 SCZ      0.0         0.0140 18.6   -0.10  -0.0307   0.0279   0.924
---
Signif. codes:  0 ‘***’ 0.001 ‘**’ 0.01 ‘*’ 0.05 ‘.’ 0.1 ‘ ’ 1
```

##### SM3 LMM for local SW triggered spindle power (Figure S5)

**Linear mixed model fit by REML**

t-tests use Satterthwaite approximations to degrees of freedom ['lmerMod']

Formula: spinpow ~ night + group + (1 | ID)

```
Analysis of Variance Table of type III  with  Satterthwaite
approximation for degrees of freedom

       Sum Sq Mean Sq NumDF  DenDF F.value   Pr(>F)
night 0.01146 0.01146     1 29.048  0.3034 0.585972
group 0.42240 0.42240     1 33.854 11.1844 0.002026 **
---
Signif. codes:  0 ‘***’ 0.001 ‘**’ 0.01 ‘*’ 0.05 ‘.’ 0.1 ‘ ’ 1
```

```
Differences of LSMEANS:
               Estimate Standard Error   DF t-value Lower CI Upper CI p-value
group HC - SCZ      0.5          0.142 33.9    3.34    0.186    0.762   0.002 **
---
Signif. codes:  0 ‘***’ 0.001 ‘**’ 0.01 ‘*’ 0.05 ‘.’ 0.1 ‘ ’ 1
```

##### SM4 Cz-O1 spindle coherence (Figure S7)

**Linear mixed model fit by REML t-tests use Satterthwaite approximations to degrees of freedom [lmerMod]**

Formula: spincoh ~ night + group + (1 | ID)

```
Analysis of Variance Table of type III  with  Satterthwaite
approximation for degrees of freedom
        Sum Sq  Mean Sq NumDF  DenDF F.value   Pr(>F)
night 0.016845 0.016845     1 30.338  3.2390 0.081857 .
group 0.041793 0.041793     1 34.705  8.0361 0.007594 **
```

##### SM5 spindle triggered Cz-O1 spindle coherence (Figure S8)

**Linear mixed model fit by REML t-tests use Satterthwaite approximations to degrees of freedom [lmerMod]**

Formula: sp_spincoh ~ night + group + (1 | ID)

```
Analysis of Variance Table of type III  with  Satterthwaite
approximation for degrees of freedom
        Sum Sq  Mean Sq NumDF  DenDF F.value   Pr(>F)
night 0.025022 0.025022     1 29.927  2.8364 0.102551
group 0.084013 0.084013     1 34.845  9.5236 0.003961 **
---
Signif. codes:  0 ‘***’ 0.001 ‘**’ 0.01 ‘*’ 0.05 ‘.’ 0.1 ‘ ’ 1
```

##### SM6 Partial least squares regression

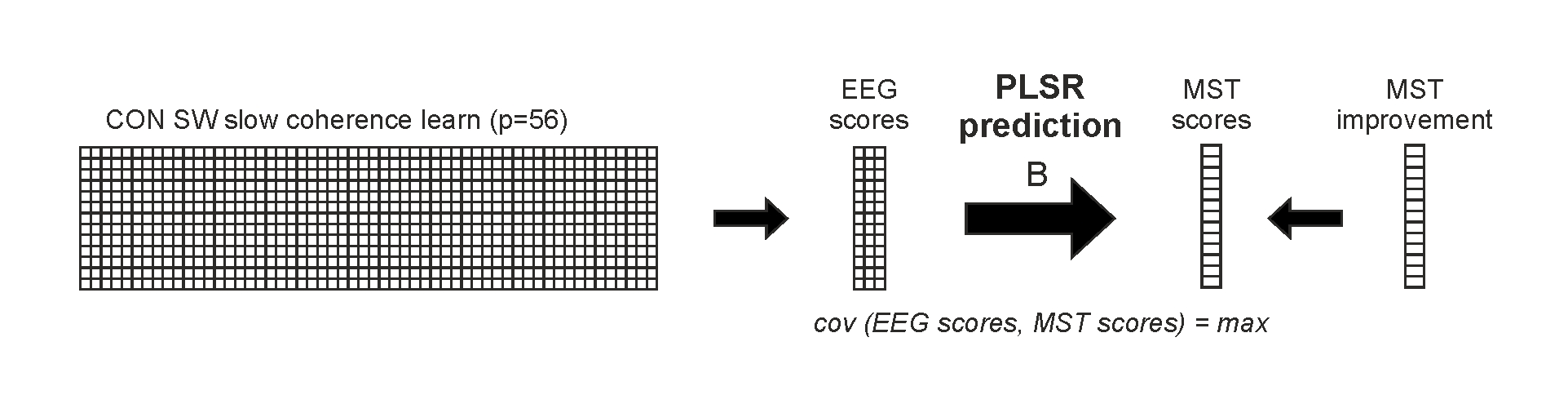

a) Partial least squares regression

PLSR seeks to maximize the covariance between linear transformations of predictor variables (EEG measures) and linear transformations of outcome variables (MST scores) in the coefficient stored in B. In this illustration we use SW triggered slow coherence values during the learning night in the control group and try to predict the outcome variable (MST change in percent) see main text methods section for more details and citations.
